# In-vivo characterization of glutamine metabolism identifies therapeutic targets in clear cell renal cell carcinoma

**DOI:** 10.1101/2022.10.31.514397

**Authors:** Akash K. Kaushik, Lindsey K. Burroughs, Amy Tarangelo, Mukundan Ragavan, Cheng-Yang Wu, Xiangyi Li, Kristen Ahumada, Vanina T. Tcheuyap, Faeze Saatchi, Quyen N Do, Cissy Yong, Tracy Rosales, Christina Stevens, Aparna Rao, Brandon Faubert, Panayotis Pachnis, Lauren G. Zacharias, Hieu Vu, Feng Cai, Thomas P. Mathews, Barbara Slusher, Payal Kapur, Xiankai Sun, Matthew Merritt, James Brugarolas, Ralph J. DeBerardinis

## Abstract

Targeting metabolic vulnerabilities has been proposed as a therapeutic strategy in renal cell carcinoma (RCC). Here, we analyzed metabolism in patient-derived xenografts (tumorgrafts) from diverse forms of RCC. Tumorgrafts from *VHL*-mutant clear cell RCC (ccRCC) retained metabolic features of human ccRCC and engage in oxidative and reductive glutamine metabolism. We used several approaches to suppress glutamine metabolism and test the effect on tumor growth. Genetic silencing of isocitrate dehydrogenase-1 or -2 impaired reductive labeling of TCA cycle intermediates and suppressed tumor growth. Glutaminase inhibition resulted in modest growth suppression and variable effects on glutamine metabolism in vivo. Infusions with [amide-^15^N]glutamine revealed persistent amidotransferase activity during glutaminase inhibition, and blocking these activities with the amidotransferase inhibitor JHU-083 also reduced tumor growth. We conclude that ccRCC tumorgrafts catabolize glutamine via multiple pathways, perhaps explaining why it has been challenging to achieve therapeutic responses in patients by inhibiting glutaminase.

**Teaser:** Glutamine fuels the TCA cycle and amidotransferase pathways in clear cell renal cell carcinoma.

## Introduction

Renal cell carcinomas (RCC) are characterized by mutations in genes that regulate intermediary metabolism, resulting in metabolic dependencies that have been proposed as therapeutic targets (*1*). Clear cell RCC (ccRCC), the most common subtype of RCC, almost universally loses the function of the von-Hippel-Lindau (*VHL*) tumor suppressor through mutation and other mechanisms(*2, 3*). The VHL protein is a subunit of an E3 ubiquitin ligase that targets the oxygen-labile α-subunits of hypoxia-inducible transcription factors (HIFs) 1 and 2 for degradation; therefore, *VHL* mutation results in sustained HIF transcriptional activity(*4*). Together, HIF-1 and HIF-2 regulate the expression of many genes, including genes involved in central carbon metabolism. HIF-1 activates the expression of glucose transporters, glycolytic enzymes, lactate dehydrogenase, and pyruvate dehydrogenase kinase-1 (*PDK1*)(*5-8*). Expression of these genes induces a phenotype of enhanced glycolysis and suppressed pyruvate oxidation in the mitochondria (*9, 10*). Suppression of pyruvate oxidation results from PDK1’s inhibitory phosphorylation of the PDH complex, which converts pyruvate to acetyl-CoA in the mitochondria (*6*).

Although the metabolic effects of *VHL* loss on metabolism are well-established in cultured ccRCC cell lines, there is a need to study these metabolic effects in tumor models directly derived from human ccRCC. It is also unclear which if any of these effects result in clinically-actionable metabolic vulnerabilities. An important bottleneck in cancer metabolism studies is the relative lack of information about pathway activity in vivo. By infusing cancer patients intra-operatively with ^13^C-labeled nutrients and examining ^13^C enrichment in metabolites extracted from the tumor after surgery, we previously characterized human tumor metabolism in several types of cancer (*11-13*). Infusion of [U-^13^C]glucose into a small cohort of ccRCC patients revealed robust ^13^C labeling of glycolytic intermediates but impaired labeling of TCA cycle intermediates in the tumor relative to adjacent kidney(*13*). This analysis revealed a metabolic phenotype consistent with the Warburg effect, as predicted by transcriptomic and metabolomic studies in ccRCC(*3, 14-17*).

It is unknown how human ccRCCs compensate for reduced pyruvate oxidation. In culture, *VHL*-mutant RCC cells use an unusual form of glutamine metabolism to supply the TCA cycle. In this pathway, termed reductive carboxylation, the reversible isoforms of isocitrate dehydrogenases (IDH1 and IDH2) catalyze the NADPH-dependent carboxylation of glutamine-derived α-ketoglutarate to produce isocitrate and citrate (*18, 19*). Citrate is then transported to the cytosol and cleaved by citrate lyase to produce acetyl-CoA, which is used to generate fatty acids, and oxaloacetate (OAA), which gives rise to other TCA cycle intermediates(*18, 19*). In ccRCC cells, reductive glutamine metabolism is a consequence of *VHL* loss because re-expressing functional *VHL* suppresses the pathway (*19, 20*). Subcutaneous xenografts derived from an established ccRCC cell line demonstrated a small amount of reductive citrate formation, raising the possibility that this pathway might contribute to central carbon metabolism in some tumors (*20*).

Identifying tumors that unequivocally require glutamine as a fuel, even from disease-relevant preclinical models, would be significant because several inhibitors of this pathway have been evaluated as therapeutic agents. Glutaminases and amidotransferases initiate glutamine catabolism by converting glutamine to glutamate. Glutaminase (GLS) is a mitochondrial enzyme that releases glutamine’s amide group as ammonia. CB-839 is a potent and well-tolerated GLS inhibitor with efficacy in some patients with heavily-pretreated RCC(*21, 22*), although recent combination studies failed to achieve primary clinical endpoints(*23*). Amidotransferases convert glutamine to glutamate, while transferring glutamine’s amide nitrogen to asparagine and intermediates in the synthesis of purines, pyrimidines, hexosamines, and nicotinamide adenine dinucleotide cofactors (*24*). The experimental amidotransferase inhibitor JHU-083, a prodrug of 6-diazo-5-oxo-L-norleucine (DON), has preclinical efficacy in multiple cancer models, suppressing tumor cell growth and enhancing antitumor immunity (*25*).

In order to identify actionable metabolic liabilities in cancer, animal models that faithfully recapitulate the metabolic features of human tumors are needed. Access to disease-relevant models is particularly important in analyzing glutamine metabolism as many cancer cell lines use glutamine as a respiratory substrate in culture (*26*), but this does not guarantee glutamine utilization in vivo. In fact, most tumors that have been carefully analyzed using isotope tracers do not use glutamine as a prominent carbon source for the TCA cycle (*27, 28*). Here, we set out to characterize the metabolic features of a large panel of patient-derived RCC tumorgrafts that retain clinical and genomic features of human tumors (*29, 30*), to determine whether they consume glutamine in vivo and respond to inhibitors of glutamine metabolism.

## Results

### *VHL*-mutant tumorgrafts recapitulate core metabolic features of primary human ccRCC

To identify tractable metabolic features in RCC, we performed metabolomics in 28 independent RCC tumorgraft lines (Table S1) passaged orthotopically in NOD-SCID mice as described previously (*29, 31*). These tumorgrafts encompass the histological and clinical diversity of aggressive forms of human RCC, including 19 high-grade ccRCCs of which 65% had confirmed *VHL* mutations, 2 each of FH-deficient RCC and papillary RCC (pRCC), 1 translocation RCC (tRCC), and 4 unclassified RCC (uRCC). Twenty-two of these tumors were derived from the primary site and the rest were from either distant (5) or regional lymph node (1) metastases. Overall, 24/28 of the models were derived from treatment-naïve tumors. Among the ccRCC tumorgrafts, 5 contained sarcomatoid features associated with poor outcomes in patients(*32*).

For metabolic analysis, we orthotopically passaged each tumorgraft into multiple NOD-SCID mice. When the tumor diameter reached ∼10-15 mm, we harvested tissues from the tumor and contralateral kidney from 2-4 mice from each model. All tissues were harvested within 3 minutes of sacrificing the mouse to minimize artifacts. We performed targeted metabolomics on 1-3 fragments from each tumor and kidney. A principal component analysis (PCA) of 118 metabolites revealed that all RCC subtypes were metabolically distinct from the contralateral kidney, but this unsupervised analysis did not distinguish RCC subtypes from each other (Fig. S1A). Supervised analysis of the 50 metabolites that best distinguished tumors from kidney revealed heterogeneity of metabolomic signatures among the ccRCC models, although most of them clustered together (Fig. S1B). As reported in ccRCC samples from patients (*16, 17, 33*), metabolites related to glutathione and glycolysis were among the most altered metabolites in tumors relative to kidneys. A supervised analysis (variable importance in the projection, VIP) identified elevated glutathione and other glutamine-related metabolites and decreased purine- and pyrimidine-related metabolites in ccRCC tumorgrafts (Fig. S1C). Similar observations have been reported in human ccRCC(*14, 16, 17*), suggesting that ccRCC tumorgrafts retain several metabolic features of human disease.

To directly determine how well the tumorgrafts recapitulate human ccRCC metabolism, we compared metabolomics data between ccRCC tumorgrafts and a large, published metabolomics dataset from human ccRCC(*17*). Unsupervised clustering of the 76 metabolites common to both datasets separated most kidney samples from tumor samples (Fig. S1D). Among the 76 metabolites detected on both platforms, 51 differed between tumor and kidney in both studies, and most of these (62%) were either accumulated or depleted in both human and mouse tumors (Fig. 1A). These commonly-perturbed metabolites included several related to central carbon and glutamine metabolism, including glutathione, 2-hydroxyglutarate, lactate, and aspartate (Fig. 1A). We performed a similar analysis between the ccRCC tumorgrafts and patient tumors from UT Southwestern’s Kidney Cancer Program (KCP) (*33*) and again observed consistent alterations in many metabolites that differentiated kidneys from tumors (Fig. S2A). This consistency is striking because although we obtained tumorgraft tissues under controlled conditions very rapidly after euthanizing the mice, we could not ensure the same consistency in the human studies. The data indicate that ccRCC tumorgrafts retain many metabolic features of human ccRCC, and that these features are durable enough to withstand multiple passages in mice.

**Figure 1.**
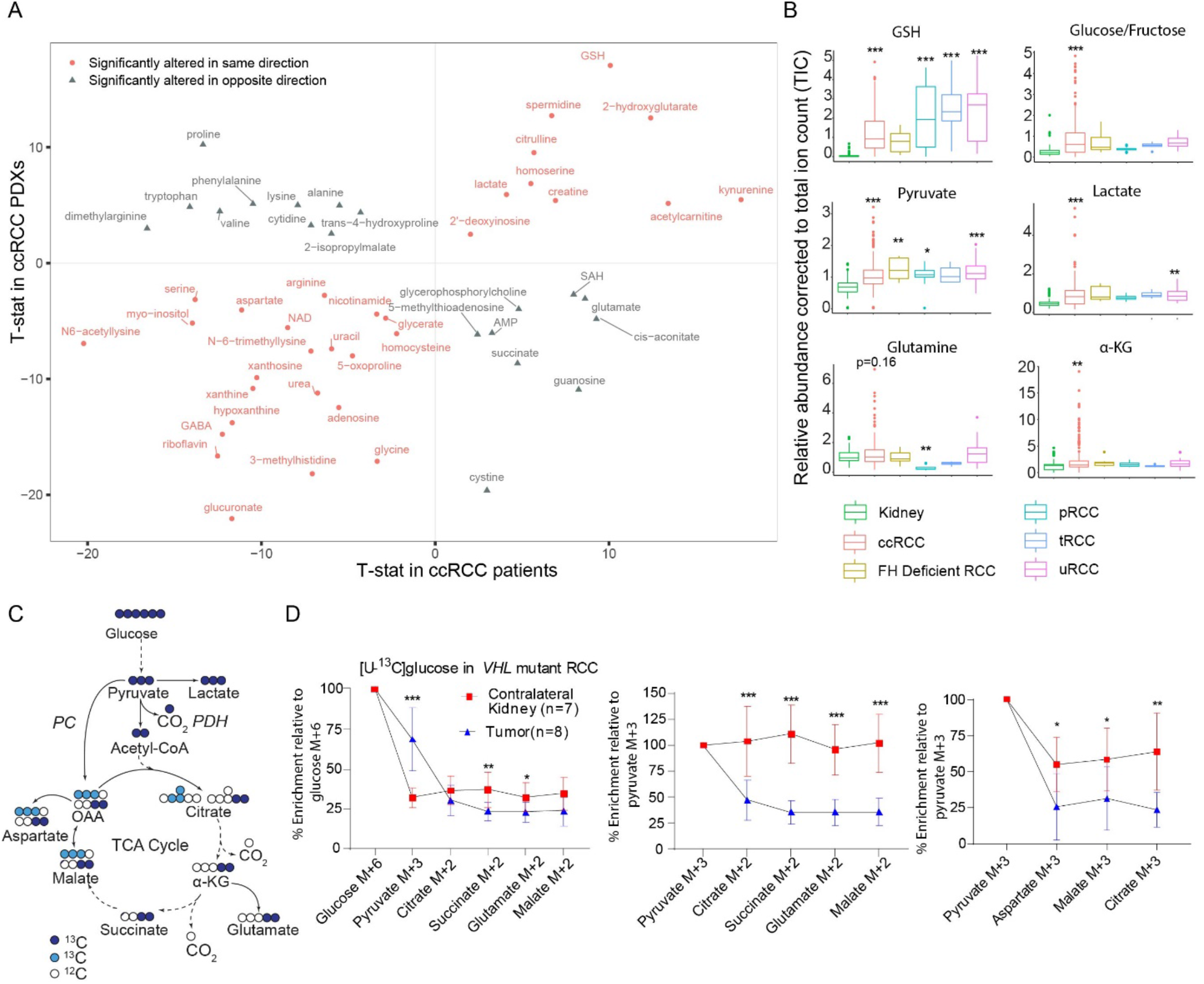
ccRCC tumorgrafts recapitulate metabolic signatures of human ccRCC. (A). Correlation plot of the metabolites significantly altered in both ccRCC tumorgrafts and human tumors. In both studies, differential metabolites between tumor and kidney were calculated using the Student’s t-test and FDR corrected (Q value<0.05). T-statistic (T-stat) was used as a surrogate for z-score to generate the correlation plot using ggplot in R. Positive T-stat values indicate metabolite elevation in tumors and negative T-stat values indicate metabolite depletion in tumors. Red circles are metabolites altered in the same direction in tumorgrafts and human ccRCC, and gray triangles are metabolites altered in opposite directions. (B). Boxplots of glutathione (GSH) and intermediates of glycolysis and the TCA cycle in RCC tumorgrafts of different histological types, including ccRCC, *FH* deficient RCC, papillary RCC (pRCC), translocation RCC (tRCC), and unclassified RCC (uRCC). The relative abundance of each metabolite was corrected to the total ion count (TIC) in each sample. One-way ANOVA coupled with pairwise t-test in R software was used to calculate statistical significance and P values were FDR corrected. (C). Illustration of central carbon flow originating with [U-^13^C]glucose. Pyruvate fuels the TCA cycle via pyruvate dehydrogenase (PDH) (dark blue circles) and carboxylases reactions such as pyruvate carboxylase (PC) (light blue circles) reactions. (D). Plots showing the percentage enrichment of metabolites relative to [U-^13^C]glucose (left panel); and the ratio of M+2 (middle panel) or M+3 (right panel) TCA cycle metabolites relative to pyruvate M+3 as surrogates of PDH and carboxylase activity, respectively. Eight mice bearing three distinct orthotopic tumorgrafts (XP258, n=3; XP374, n=3; XP490, n=2) were infused with [U-^13^C]glucose for 3 hours, and metabolites extracted from tissues were analyzed using GC-MS. M is the mass of the unlabeled metabolite. P-values were calculated between all tumors and kidney using the Student’s t-test. P values: ***<0.001, **<0.01, *<0.05

Next, we compared metabolomic alterations among the different RCC subtypes represented in the tumorgraft panel. Glutathione was markedly elevated in all subtypes relative to kidney (Fig. 1B). Glucose/fructose (our platform did not distinguish these from each other), lactate, and pyruvate from the glycolytic pathway were elevated in tumorgrafts derived from primary ccRCCs, but only inconsistently in tumorgrafts from metastatic ccRCC and other RCC subtypes (Fig. 1B). Levels of several TCA cycle and related metabolites were also altered in the ccRCC tumorgrafts, with an enhanced abundance of α-ketoglutarate and 2-hydroxyglutarate and a trend towards elevated glutamine and decreased succinate (Fig. 1B, Fig. S2B), as reported in human ccRCC (*17, 33*).

### Orthotopic *VHL*-mutant tumorgrafts display reduced pyruvate oxidation relative to the non-malignant kidney

We previously showed using isotope labeled glucose infusions in patients that human ccRCCs display reduced pyruvate oxidation compared to patient-matched kidney tissue (*13*). To test whether the same phenotype is observed in ccRCC tumorgrafts, we intravenously infused [U-^13^C]glucose into NOD-SCID mice with orthotopic *VHL*-mutant ccRCC. This technique assesses the fates of carbon from plasma glucose into glucose-dependent pathways in the tissues, including glycolysis and the TCA cycle (Fig. 1C). The primary ccRCC models displayed elevated ^13^C labeling in pyruvate compared to the contralateral kidney but reduced labeling in TCA cycle-related metabolites (Fig. 1D). M+2 labeling in the TCA cycle intermediates relative to pyruvate m+3 can be used as a surrogate for carbon entry through pyruvate dehydrogenase (PDH), and m+3 labeling in the TCA cycle intermediates relative to pyruvate m+3 can be used as a surrogate for carbon entry via carboxylases, e.g. by pyruvate carboxylase (PC) (Fig. 1C). The kidneys displayed good propagation of labeling from pyruvate into the TCA cycle via both pathways, but these ratios were suppressed in the tumors (Fig. 1D). This is similar to the labeling pattern observed in ccRCC patients infused with [U-^13^C]glucose during nephrectomy (*13*). Overall, in both metabolomics and isotope tracing assays, these tumorgrafts recapitulate metabolic features of human ccRCC and provide an opportunity to identify nutrients used by these tumors.

### Glutamine fuels the TCA cycle in *VHL-*mutant RCC

The elevation of glutamine-related metabolites and suppressed contribution of glucose to the TCA cycle suggested that glutamine is a carbon source for the TCA cycle in ccRCC tumorgrafts. To test this, we infused [U-^13^C]glutamine into mice bearing five independent *VHL*-mutant ccRCC tumorgrafts with genetic and histological heterogeneity (Fig. S3A). We used an infusion method that labels 30-40% of glutamine and 10-17% of glutamate in the plasma (Fig. S3B). Glutamine enrichment did not differ between the tumors and adjacent benign or contralateral kidney (Fig. S3C). However, 4/5 tumorgrafts had elevated labeling in glutamate relative to the plasma and non-malignant kidney, suggesting enhanced conversion of glutamine to glutamate in the tumors (Fig. 2A, Fig. S3B). Glutaminase (GLS) was expressed in all tumorgrafts at levels comparable to the kidney (Fig. S3D).

**Figure 2.**
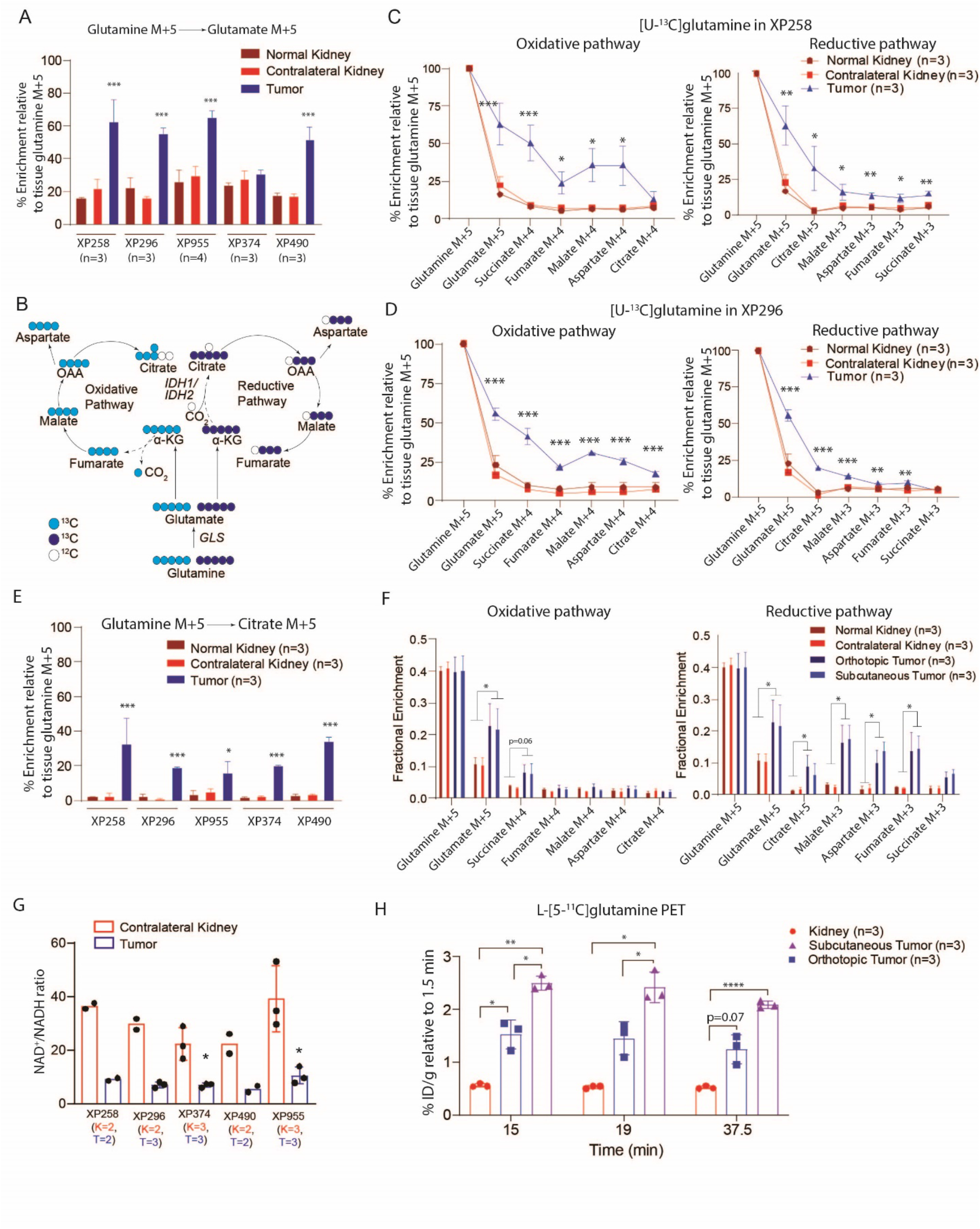
Glutamine is a carbon source for the TCA cycle in ccRCC tumorgrafts. (A). Percentage enrichment of glutamate M+5 relative to glutamine M+5 in orthotopic tumors, normal kidney, and contralateral kidney in mice infused with [U-^13^C]glutamine. One-way ANOVA was used to assess the statistical significance of glutamate M+5 enrichment. (B). Schematic of the carbon flow from [U-^13^C]glutamine into the TCA cycle via the oxidative and reductive pathways represented by light and dark blue circles, respectively. In the first cycle of the oxidative pathway, four carbons of the TCA cycle intermediates become labeled with ^13^C. Citrate is labeled with five ^13^C in the first cycle of the reductive pathway, with subsequent generation of OAA, malate and fumarate containing three ^13^C labels. (C). Percentage enrichment of TCA cycle intermediates relative to [U-^13^C]glutamine in XP258 tumorgrafts, adjacent benign kidney (labeled as a normal kidney), and contralateral kidney. The left panel shows ^13^C labeling via the oxidative pathway and the right panel represents the ^13^C labeling via the reductive pathway. The experiment was conducted in a minimum of 3 mice per tumorgraft. One-way ANOVA coupled with a pairwise t-test was used to assess the statistical significance of ^13^C enrichment between tissues. (D). Similar experiment as in panel C, but in XP296 tumorgrafts. (E). Percentage enrichment of citrate M+5 relative to [U-^13^C]glutamine (glutamine M+5) across all tumorgraft and kindey tissues. The experiment was conducted in a minimum of 3 mice per tumorgraft model. One-way ANOVA coupled with a pairwise t-test was used to assess the statistical significance of ^13^C enrichment in citrate M+5 between tissues. (F). Percentage enrichment of ^13^C relative to [U-^13^C]glutamine in the intermediates of TCA cycle via both oxidative (left panel) and reductive (right panel) metabolism in orthotopic tumors, subcutaneous tumors, normal kidney, and contralateral kidney. Data are plotted from experiments conducted in three mice implanted with both orthotopic and subcutaneous XP955 tumorgrafts. One-way ANOVA was used to assess the statistical significance of ^13^C enrichment. (G). NAD^+^/NADH ratio in ccRCC tumorgraft and contralateral kidney tissues. The number of tissues (n) used for analysis is displayed below each plot as K=n for kidney and T=n for tumor. The Student’s t-test was used to assess the significance. (H).^11^C signal relative to 1.5 min (time of maximal signal in the kidney) in kidney and tumors. We used one-way ANOVA to determine the statistical significance. P values: ****<0.0001,***<0.001, **<0.01, *<0.05

Next, we examined the contributions of glutamine-derived carbon into TCA cycle metabolites, assessing both the oxidative and reductive pathways of glutamine metabolism (Fig. 2B). [U-^13^C]glutamine labels glutamate and α-KG as m+5. Oxidation of α-KG results in m+4 labeling in other TCA cycle intermediates, while reductive carboxylation generates m+5 citrate and m+3 in other metabolites(*18*). Most of the tumors had elevated ^13^C labeling in α-KG compared with kidney tissues (Fig. S3E). All tumors displayed elevated ^13^C labeling in at least some of the other TCA cycle intermediates as well. XP258 and XP296 demonstrated enhanced labeling along both pathways, with reductive labeling particularly evident in citrate and oxidative labeling characterizing the rest of the cycle (Fig. 2C, D). In both of these models, oxidative labeling exceeded reductive labeling in all metabolites besides citrate, indicating flow around the TCA cycle to OAA, but suppressed formation of citrate via citrate synthase, likely because of reduced PDH activity (*34, 35*). XP955 demonstrated more prominent reductive labeling and the other two lines had less consistent labeling around the cycle (Fig. S2F-H). All five tumors had enhanced citrate m+5 (reductive) labeling relative to the kidney, even XP374, which did not display elevated glutamate labeling relative to the kidney (Fig. 2E). The increased contribution of glutamine is notable in that glutamine catabolism in tumors analyzed by ^13^C tracing in vivo is generally low (*27, 28*).

To confirm the reductive formation of citrate in these tumors, we infused [1-^13^C]glutamine. In this labeling scheme, ^13^C is released by α-KG dehydrogenase but retained in citrate when α-KG becomes carboxylated (Fig. S4A). *VHL*-mutant tumorgrafts (XP258 and XP955) contained enhanced citrate m+1 relative to healthy kidney tissues (Fig. S4A).

Because the microenvironment influences tumor metabolism (*36, 37*), we asked whether the site of implantation impacts glutamine-dependent labeling. XP955 tumors were implanted into the subcutaneous space and the renal capsules in the same mice. Then the mice were infused with [U-^13^C]glutamine and metabolites were extracted from tumors at both sites. Absolute ^13^C enrichment in glutamate and TCA cycle intermediates were nearly identical between subcutaneous and orthotopic tumors, with tumors at both sites demonstrating increased labeling in TCA cycle intermediates relative to non-malignant kidney (Fig. 2F). This experiment implies that glutamine handling in these tumors results from cancer cell-autonomous properties rather than the microenvironment.

Compared with the other *VHL*-mutant tumors, XP955 tumors had the highest labeling in aspartate and malate from the reductive pathway, and M+3 labeling in these metabolites exceeded M+5 labeled citrate (Fig. S3G). This is an unusual pattern because citrate is upstream of aspartate and malate in this pathway and would presumably have equivalent or higher labeling (Fig. 2 B). To provide further detail to the tracer analysis, we used NMR to examine the ^13^C positional enrichment in metabolites after infusion with [U-^13^C]glutamine. Glutamine oxidation produces uniformly-labeled malate on the first turn of the TCA cycle, so spin-coupling generates complex NMR multiplets at malate C2 and C3. Specifically, oxidative metabolism should produce quartets (i.e., doublets of doublets), reflecting labeling in C1-3 observable at the C2 chemical shift and C2-4 observable at the C3 chemical shift. Reductive metabolism generates citrate labeled at all positions except C6, and cleavage by ATP citrate lyase generates OAA and malate labeled in C2-4 (Fig. S4B). Equilibration with fumarate, a symmetric metabolite, ultimately results in malate labeled in either C1-3 or C2-4 in equal measures. Therefore, reductive metabolism also produces multiplets at malate C2 and C3, including quartets. However, unlike the uniformly labeled malate from the oxidative pathway, malate from the reductive pathway lacks ^13^C on either C1 or C4. This produces malate C1-3 and C2-4 isotopomers that give rise to doublets at the C2 and C3 positions in addition to contributions from the quartets (i.e., at C2, the doublets reflect malate molecules containing ^13^C at C2 and C3 but not C1; and at C3, malate molecules containing ^13^C at C2 and C3 but not C4). These doublets should be similar in size at steady state.

NMR spectra of metabolites extracted from [U-^13^C]glutamine-infused tumors revealed complex C2 and C3 malate multiplets, but little malate signal was detected in spectra from the kidney (Fig. S4C, D). In tumor metabolites, quartets and 2-3 doublets were prominent at both C2 (C2D23) and C3 (C3D23), consistent with reductive labeling in agreement with the mass spectrometry data. To more directly assess the contribution of oxidative metabolism, we examined 1-2 and 3-4 doublets in malate; these patterns arise on the second oxidative turn of the cycle and were barely detectable (Fig. S4B-D). Some ^13^C label was also detected at resonances for C2/C4 in tumor-derived citrate (Fig. S4C, E), but the signal-to-noise ratio was much lower than for malate, again consistent with the low citrate labeling by mass spectrometry. Altogether, these data indicate prominent reductive and suppressed oxidative labeling from glutamine in this ccRCC model. The low citrate enrichment may reflect contributions of an unlabeled citrate pool arising from sources other than glutamine. However, glutamine-derived citrate contributes to other TCA cycle intermediates, resulting in relatively high enrichment in these metabolites.

Reductive carboxylation in cultured cells has been ascribed to an increased α-KG/citrate ratio or a shift in the cellular redox ratio to a more reduced state (i.e., decreased NAD^+^/NADH ratio) (*20, 35*). NAD^+^ and NADH measurements in these tumors revealed lower NAD^+^/NADH ratios than in non-malignant kidneys (Fig. 2G, Fig. S5A). Although both α-KG and citrate levels tended to be elevated in tumors, the α-KG/citrate ratio was not markedly different from the kidneys (Fig. S5B).

ccRCC tumorgrafts contained elevated levels of GSH and 2-hydroxyglutarate (Fig. 1B, Fig. S2B), both of which are also abundant in human ccRCC^(*14, 17, 20, 33, 38*)^. Infusion with [U-^13^C]glutamine resulted in labeling in both metabolites (Fig. S5C, D). In GSH, the fractional enrichment was similarly low in the tumor and kidney, but the accumulation of GSH in the tumors translated into a much higher abundance of labeled GSH. Labeled 2-HG was present in the tumors but not the kidneys and was nearly all L-2HG (Fig. S5D, E), as previously reported in the human ccRCC (*38*).

### L-[5-^11^C]glutamine positron emission tomography and computed tomography (PET/CT) in ccRCC

To assess glutamine metabolism using a different method, we performed PET/CT with L-[5-^11^C]glutamine in mice bearing XP490 tumorgrafts in orthotopic and subcutaneous regions. Given the low soft-tissue contrast of CT, we assessed the anatomical location of the tumors and contralateral kidney with MRI (Fig. S6A). The dynamic distribution of L-[5-^11^C]glutamine should reflect the metabolism (uptake and retention) of glutamine by the tumors as long as the ^11^C label remains in the tissue. In the kidney, the time-activity curve of ^11^C radioactivity (percent injected dose per gram, %ID/g) reached its maximum (∼ 20

%ID/g) within 1.5 minutes of injection and then dropped to about half after 37.5 minutes, indicating a significant washout of L-[5-^11^C]glutamine. This observation was similarly reported in patients with metastatic colorectal cancer (*39*). In the tumors, although the peak PET signal was lower than that in kidney, it was retained over the entire time course (Fig. S6B). A significantly increased ratio of signal uptake relative to the peak time point demonstrated higher signal retention in the tumors (both orthotopic and subcutaneous) relative to kidney (Fig. 2H). This retention is consistent with glutamine’s extensive participation in metabolism in these tumors.

### IDH1 and IDH2 support ccRCC growth in vivo

Reductive carboxylation is catalyzed by NADPH-dependent cytosolic IDH1 and mitochondrial IDH2 (Fig. S7A) (*40*). To study the mechanism of reductive carboxylation in vivo and the role of IDH1 and IDH2 in tumorgraft growth, we generated a cell line from the *VHL*-mutant tumorgraft XP258. Like XP258 tumorgrafts, this cell line displayed both oxidative and reductive metabolism of glutamine in culture, and xenografts generated from XP258 cells after several passages in culture displayed nearly identical patterns of glutamine metabolism as the original tumorgrafts (Fig. 3A).

**Figure 3.**
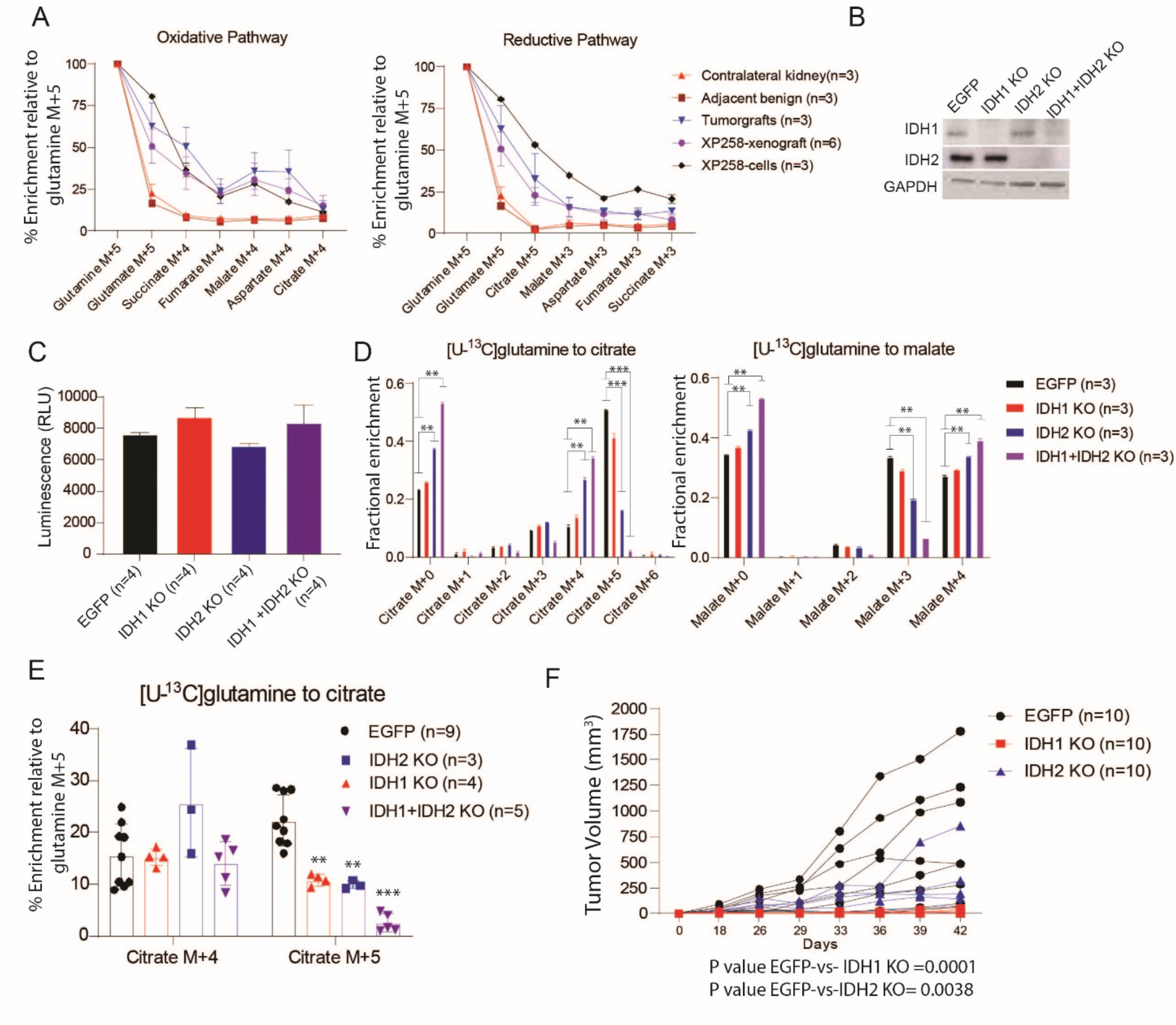
IDH1 and IDH2 regulate reductive carboxylation and ccRCC growth. (A). Comparison of the percentage enrichment of ^13^C relative to [U-^13^C]glutamine in the intermediates of TCA cycle between XP258 orthotopic PDX, a cell line derived from this PDX (XP258-cells), and subcutaneous xenografts generated from the cell line (XP258-xenografts). Tumor-bearing mice were infused with [U-^13^C]glutamine for 5 hours and XP258 cells were labeled in culture with [U-^13^C]glutamine for 4 hours. The left panel corresponds to ^13^C labeling in the oxidative pathway and the right panel corresponds to ^13^C labeling in the reductive pathway. (B). Western blot shows abundance of IDH1 and IDH2 in single and double KO (IDH1 + IDH2 KO) cells generated using lentiviral CRISPR-Cas9. EGFP is a control cell line with a gRNA targeting a non-human, GFP (green fluorescence protein) expressing gene. (C). Plot showing cell viability of EGFP, IDH1 KO, IDH2 KO, and IDH1 +IDH2 KO XP258 cells assessed using Promega CellTiter-Glo® assay. (D). Fractional enrichment of ^13^C labeled isotopologues of citrate and malate from EGFP, IDH1 KO, IDH2 KO, and IDH1 KO +IDH2 KO (double KO) cells after 4 hours of culture with [U-^13^C]glutamine. One-way ANOVA was used to assess the statistical significance of ^13^C enrichment in citrate and malate isotopologues between each cell line (n=3). One-way ANOVA with a pairwise t-test was used to assess the statistical significance. (E). Percentage enrichment of citrate M+4 and citrate M+5 relative to [U-^13^C]glutamine in EGFP, IDH1 KO, IDH2 KO, and double KO (IDH1 KO+ IDH2 KO) XP258 tumors. Mice were infused with [U-^13^C]glutamine for 5 hours before tissues were harvested. Data are represented from three or more xenografts generated subcutaneously in NOD-SCID mice using the above cell line. One-way ANOVA with a pairwise t-test was used to determine the statistical significance. (F). Growth of EGFP, IDH1 KO, and IDH2 KO subcutaneous XP258 xenografts in NOD-SCID mice (n=10, each cell line). Tumors were measured periodically using calipers. P values were calculated using a mixed effect two-way ANOVA that assessed the significance of tumor growth with time between EGFP vs. IDH1 KO and EGFP vs. IDH2 KO tumors. P values: ***<0.001, **<0.01, *<0.05

We next used CRISPR-Cas9 to create pools of cells lacking IDH1, IDH2, or both (Fig. 3B). Cells lacking IDH1, IDH2, or both proliferated in the culture at the same rate as parental cells (Fig. 3C). Reductive labeling of citrate from [U-^13^C]glutamine was only marginally changed by IDH1 loss, but IDH2 loss had a more substantial effect, and reductive labeling was nearly eliminated in cells lacking both IDH1 and IDH2 (Fig. 3D). In cells lacking IDH2 or both IDH1 and IDH2, a higher fraction of citrate and malate displayed labeling through the oxidative pathway (Fig. 3D). Loss of IDH2 or both IDH1 and IDH2 also resulted in enhanced labeling of TCA cycle intermediates from [U-^13^C]glucose (Fig. S7B).

We next studied the role of IDH1 and IDH2 in xenograft metabolism and growth. Xenografts from either IDH1 KO or IDH2 KO had a reduced fraction of citrate formed through reductive carboxylation, and this fraction was nearly eliminated by loss of both enzymes (Fig. 3E). The loss of IDH1 or IDH2 also reduced tumor growth (Fig. 3F). We could not consistently generate enough palpable tumors from cells lacking both IDH1 and IDH2 to perform xenograft growth experiments, despite their normal proliferation rate in culture. Examination of protein expression at the end of these experiments revealed that most tumors re-expressed some IDH1 and IDH2 (Fig. S7C). We confirmed this in a second cell line generated from XP258 tumorgrafts, again noting xenograft growth suppression but some recovery of IDH1 and IDH2 expression by the end of the experiment (Fig. S7D-F). Altogether, the data indicate that IDH1 and IDH2 are dispensable in culture, even in cells that display significant levels of reductive metabolism. However, both enzymes contribute to XP258 tumor growth in vivo.

### The glutaminase inhibitor CB-839 moderately decreases glutamine metabolism and growth in ccRCC tumorgrafts

We next examined the effects of CB-839 on tumor metabolism and growth. We generated subcutaneous tumors from XP490, XP258, and XP296 tumorgrafts. XP490 and XP258 were generated from treatment-naïve patients, and XP296 was generated from a patient previously treated with everolimus. CB-839 decreased glutamine’s contribution to the TCA cycle along both the oxidative and reductive pathways in XP296 and XP490, but not in XP258 tumors (Fig. 4A, C, E). CB-839 also decreased glutamine’s contribution to glutathione and L-2HG in XP296 tumors (Fig. S8A, B). Despite the variable impact on glutamine metabolism, all three tumorgrafts displayed modest growth suppression upon CB-839 treatment, with most tumors displaying stable tumor volume over three weeks of treatment (Fig. 4B, D, F). Tumor regression was not observed. Body weights were largely unaffected by CB-839 (Fig. S8C).

**Figure 4.**
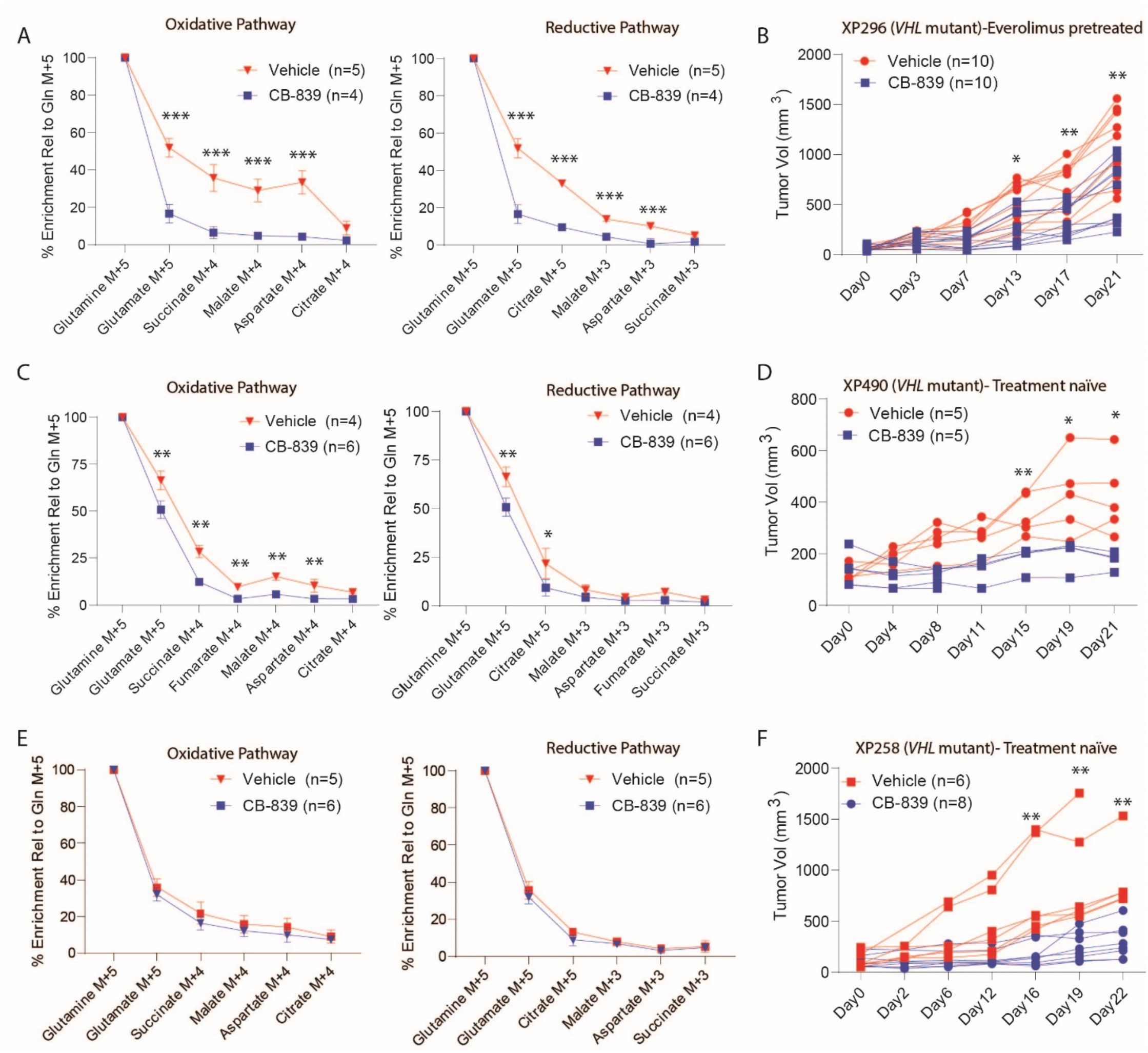
CB-839 variably decreases glutamine metabolism and ccRCC growth. (A). Percentage enrichment of ^13^C labeled TCA cycle intermediates relative to [U-^13^C]glutamine in the oxidative (left panel) and reductive (right panel) pathway in XP296 tumorgrafts bearing mice treated with 200 mg/kg of CB-839 or vehicle for 21 days. Student’s t-test was used to assess the statistical significance of ^13^C enrichment between CB-839 and vehicle-treated tumors (n≥4). (B). Spider plot of growth of subcutaneous XP296 tumorgrafts in NOD-SCID mice treated twice daily with 200 mg/kg of CB-839 or vehicle. Treatment was started when tumors reached 100-200 mm^3^. Data are plotted from two independent experiments. The Student’s t-test was used to calculate the statistical significance of tumor growth at each time point (n=10, each treatment arm). (C). Similar to A, but for XP490 tumorgrafts. (D). Similar to B, but for XP490 tumorgrafts. (E). Similar to A, but for XP258 tumorgrafts. (F). Similar to B, but for XP258 tumorgrafts. P values: ***<0.001, **<0.01, *<0.05

### Amidotransferases inhibitor JHU-083 decreases glutamine metabolism and tumor growth in *VHL* mutant ccRCC

Despite the large effect of CB-839 on carbon transfer from glutamine into TCA cycle intermediates in XP296 and XP490, residual glutamate labeling was noted in all tumors (Fig. 4A, C, E). This was not unexpected, as amidotransferases and other enzymes convert glutamine to glutamate and might compensate for glutaminase blockade (Fig. S8D) (*24, 41*). To examine the contribution of these reactions, we infused glutamine labeled on the amide nitrogen, [amide-^15^N]glutamine, into mice with XP296, XP258 and XP490 tumorgrafts, and examined isotope enrichment in metabolites in the presence and absence of CB839. We first assessed labeling in orotate, an intermediate in pyrimidine metabolism that acquires nitrogen from the amide position of glutamine through activity of the trifunctional CAD (carbamoyl-phosphate synthetase-2/aspartate transcarbamylase/dihydroorotase) enzyme. Orotate was highly labeled in all tumors studied and enrichment tended to be higher in the tumors than the blood, consistent with local production (Fig. S8E-G). Orotate was also labeled in tumors treated with CB839 (Fig. S8E-G), indicating persistent amidotransferase activity during glutaminase inhibition.

Other metabolites were also labeled from [amide-^15^N]glutamine. Citrulline, a urea cycle intermediate that acquires nitrogen from free ammonia via the carbamoyl-phosphate synthetase-1 and ornithine transcarbamylase activities in liver mitochondria, was also highly labeled. However, citrulline labeling was equivalent between tumor and plasma and was suppressed by CB-839 (Fig. S8E-G). These findings suggest that unlike orotate, tumors acquire labeled citrulline primarily from the blood. CB-839 may suppress citrulline labeling because ammonia generated through glutaminase activity in the liver and elsewhere supplies the urea cycle.

We next studied the effect of amidotransferase inhibition on glutamine metabolism and growth in ccRCC xenografts. Mice with XP258 and XP296 tumors were treated with JHU-083. Infusions with [U-^13^C, ^15^N]glutamine were performed in other mice earlier on treatment to determine the metabolic effects of JHU-083. Several metabolites from pathways involving amidotransferases, including asparagine, guanosine and cytosine monophosphate, displayed suppressed labeling upon JHU-083 treatment (Fig. 5A). However, ^13^C labeling of TCA cycle intermediates was either unaffected or increased by JHU-083 (Fig. 5B). Labeling of these metabolites was suppressed by CB-839, suggesting that JHU-083 inhibits amidotransferases but not glutaminase at these doses in this model. Despite the persistence of glutaminase activity, JHU-083 suppressed the growth of XP258 and XP296 tumors (Fig. 5C).

**Figure 5.**
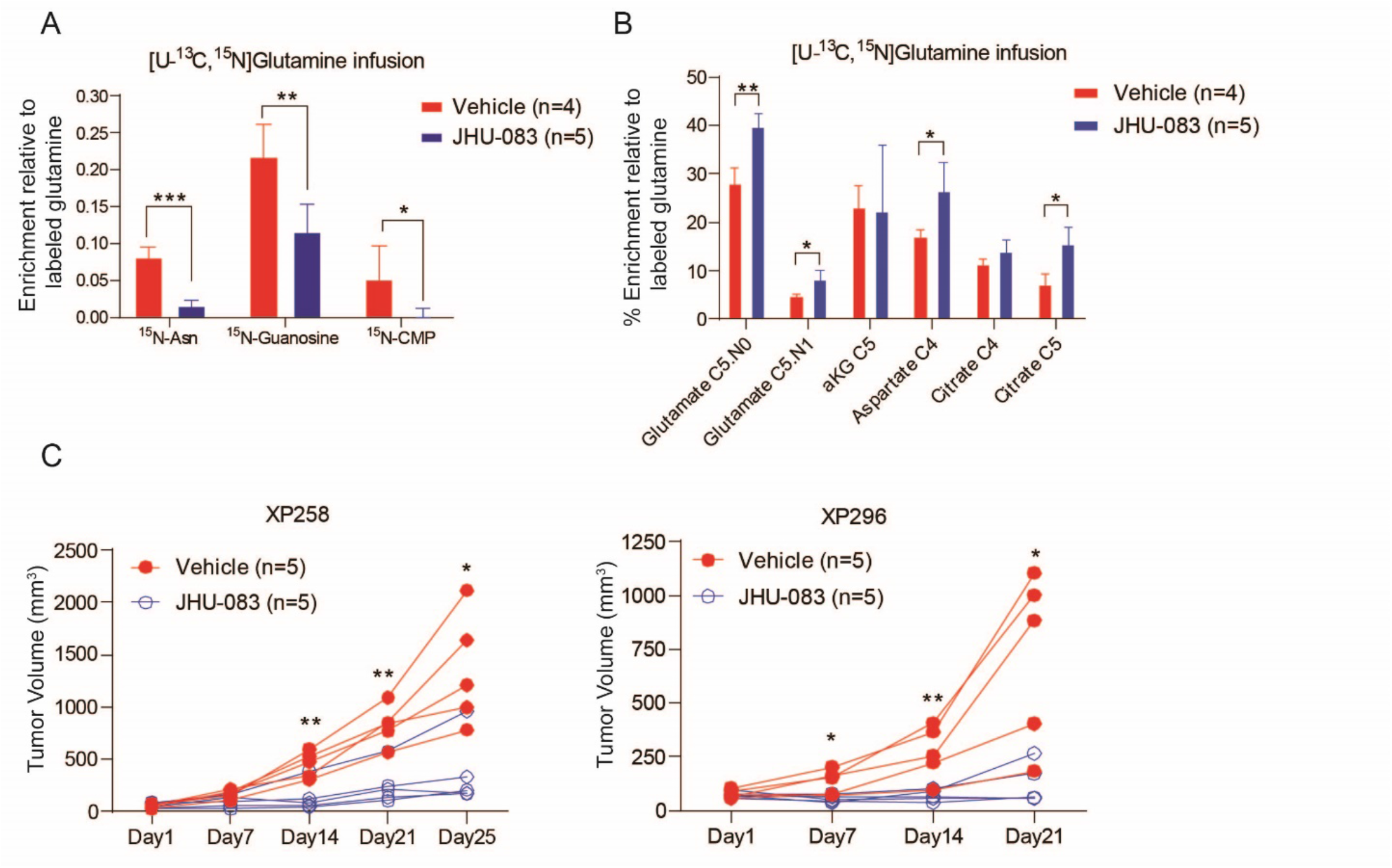
JHU-083 decreases glutamine-dependent nitrogen metabolism and ccRCC growth. (A). Percentage enrichment of ^15^N labeled asparagine (Asn), guanosine, and cytidine monophosphate (CMP) relative to [U-^13^C,^15^N]glutamine in XP258 tumorgrafts treated with 1.83 mg/kg of JHU-083 or vehicle for 5 days. The Student’s t-test was used to assess the statistical significance of ^15^N enrichment between JHU-083 and vehicle-treated tumors (n≥4). (B). Percentage enrichment of ^13^C labeled TCA cycle intermediates relative to [U-^13^C, ^15^N]glutamine in XP258 tumorgrafts treated with JHU-083 and vehicle in panel a. The Student’s t-test was used to assess the statistical significance of ^13^C enrichment between JHU-083 and vehicle-treated tumors (n≥4). (C). Spider plot of growth of subcutaneous XP258 tumorgrafts in NOD-SCID mice treated with 1.83 mg/kg of JHU-083 or vehicle for 25 days (5 days on and two days off treatment). Treatment was started when tumors reached ∼50 mm^3^. The Student’s t-test was used to calculate the statistical significance at each time tumor growth was measured (n=5, each treatment arm). (D). Same as c, but for XP296 tumorgrafts treated with 1.83 mg/kg of JHU-083 and vehicle for 21 days. P values: ***<0.001, **<0.01, *<0.05

## Discussion

Metabolic phenotypes in human tumors are incompletely defined, and we seldom know how well key metabolic features in human tumors are recapitulated in mice. This presents challenges in using mouse models to identify metabolic liabilities in human tumors. Human ccRCC metabolism has been well characterized through metabolomics and isotope labeling studies, which have emphasized enhanced glycolysis and suppressed pyruvate oxidation (the Warburg effect), features also observed in *VHL*-deficient ccRCC cell lines(*13*),(*14, 16, 17*). We extended these comparisons here by characterizing metabolism in a large series of orthotopic patient-derived tumorgrafts that retain genomic and histological features of human RCC (*30*). ccRCC-derived tumorgrafts display many metabolic features of human ccRCC, including suppressed contributions of glucose to TCA cycle intermediates and accumulation of glutamine-derived metabolites such as glutathione and L-2HG. We find from two kinds of tracer studies, [^13^C]glutamine infusion and dynamic [^11^C]glutamine PET, that glutamine is a carbon source for intermediary metabolism in these tumors, with enhanced contributions relative to nonmalignant kidney. The PET data indicate retention of glutamine carbon (specifically, carbon 5) within the tumors. Whether this primarily reflects glutamine’s contribution to the TCA cycle, or to glutathione, proteins and other macromolecules, is unknown. We also note that glutamine metabolism is an intrinsic characteristic of these tumors rather than simply a consequence of the microenvironment, as both ^11^C and ^13^C labeling features were similar between orthotopic and subcutaneous sites. A recent study that used radiotracers to compare glutamine uptake between cancer cells and immune cells in the tumor microenvironment also concluded that glutamine is preferentially taken up and retained by malignant cells (*42*).

Our ^13^C labeling approach reports fractional enrichment of metabolites downstream of the tracer after several hours of exposure to ^13^C-glutamine. Therefore, a limitation of the approach is that it reports glutamine’s metabolic fates but not the rates of glutamine-dependent pathways, including glutamine oxidation and reductive carboxylation. Nevertheless, glutamine’s prominent contributions to several pathways in the tumors compared to the kidneys raised the possibility of vulnerabilities associated with these pathways. We confirmed this by blocking IDH1/2, glutaminase and amidotransferases. Combining these loss-of-function experiments with isotope tracing is informative because it provides direct evidence that the target enzyme contributes to the pathway in vivo.

Numerous studies in cultured cells have demonstrated enhanced reductive labeling of TCA cycle intermediates under conditions of *VHL* loss, hypoxia, PDH suppression, or defects in the TCA cycle or electron transport chain (ETC) (*18, 19, 34, 35*). A small amount of reductive carboxylation was also observed in xenografts of *VHL*-deficient UMCR3 cells, and growth of these tumors was suppressed by BPTES, a preclinical GLS inhibitor (*20*). The tumors in the current study generally show combined oxidative and reductive labeling, with the latter resulting in citrate m+5 during infusion with [U-^13^C]glutamine. This phenomenon may be related to PDH suppression, as a reduced citrate:α-KG or NAD^+^/NADH ratio increases the appearance of reductively labeled citrate (*20, 35*). We do not know whether these tumorgrafts contain a net reductive flux from α-KG to citrate, but the presence of citrate m+5 prompted us to examine the role of IDH1 and IDH2. Both IDH1 and IDH2 contribute to citrate m+5 labeling in vivo. More importantly, despite the fact that these enzymes are dispensable for cell growth in culture, the loss of either one reduces tumor growth in vivo. Difficulties establishing tumors lacking both IDH1 and IDH2, and the eventual re-expression of IDH1 and IDH2 during tumor growth, implies that these enzymes provide necessary metabolic functions in vivo. At present, we do not know if reductive citrate formation per se is the key IDH1/2 function in vivo, but the data suggest that dual targeting of these enzymes could produce a substantial therapeutic effect. A compound that inhibits the oncogenic, mutant forms of both IDH1 and IDH2 has been developed (*43*), so perhaps potent inhibition of both wild-type enzymes could be achieved with a single compound.

Because glutamine is a prominent carbon and nitrogen source, there has been a sustained interest in suppressing glutamine catabolism in tumors. Several recent studies have shown preclinical efficacy of targeting glutamine metabolism in vivo by inhibitors of glutamine transporters, glutaminase, and amidotransferases (*25, 41, 44-48*). Phase-1 dose-escalation clinical studies with CB-839 demonstrated good tolerability of the drug, and stable disease or partial response in some RCC patients (*22*). However, the CANTATA (NCT03428217) Phase 3 trial revealed that combining CB-839 with cabozantinib in advanced, metastatic RCC patients pre-treated with checkpoint inhibitors or anti-angiogenic therapies did not improve outcomes over cabozantinib alone (*49*). Our data provide some insight into how tumors may resist CB-839. Upon treatment with CB-839, ccRCC tumorgrafts derived from treatment-naïve primary tumors display modest growth suppression and metabolic derangements. All tumors contained ^13^C-labeled glutamate and downstream metabolites despite CB-839 therapy, indicating persistent glutamine metabolism in the tumors. Glutamate import from the circulation could also provide a residual source of glutamine-derived glutamate, but labeling from the circulating ^13^C-glutamate pool in these experiments was less than 5% (data not shown). Transfer of ^15^N from [amide-^15^N]glutamine or [U-^13^C,^15^N]glutamine to products of amidotransferase reactions, including asparagine and intermediates from nucleotide metabolism, indicates that this family of enzymes enables residual glutamine catabolism in CB-839-treated tumors.

Since the early 1950s, clinical studies in several kinds of cancer observed stabilization or remission in some patients treated with low-dose DON (*50, 51*). However, clinical development was halted due to inconsistent response and gastrointestinal toxicity in dose-escalation studies (*52*). The recent development of JHU-083, a chemically-modified prodrug of DON with selective activation in the tumor microenvironment and a reduced toxicity index, has suggested an alternative approach to broad inhibition of glutamine catabolism in cancer. In syngeneic tumors, JHU-083 impairs glutamine catabolism in cancer cells and produces a nutrient microenvironment more amenable to the antitumor effects of cytotoxic T cells and other immune cells (*25, 53, 54*). JHU-083’s efficacy in the immunodeficient tumorgraft models described here may underestimate its overall efficacy in tumors with robust glutamine catabolism. This is particularly relevant for ccRCC, a highly immunogenic tumor type with clinical responses to immunotherapy (*55, 56*). Our isotope-tracing experiments with [U-^13^C, ^15^N]glutamine showed decreased ^15^N-enrichment in asparagine and nucleotide intermediates upon JHU-083 treatment. Surprisingly, JHU-083 did not reduce ^15^N-orortate enrichment in the tumors (data not shown). This may indicate that ^15^N-labeled orotate is imported from the circulation, or that some of the ^15^N in orotate is derived from ^15^N-aspartate synthesized elsewhere in the mouse. Importantly, although several studies have reported that JHU-083 decreases glutaminase activity in tumors (*53, 57*), the effect appears to be different in ccRCC tumorgrafts. In our models, ^13^C labeling of TCA cycle intermediates is glutaminase-dependent and inhibited by CB-839. As expected, this drug does not impair amidotransferase reactions. On the other hand, JHU-083 inhibits amidotransferase reactions but not ^13^C labeling of TCA cycle intermediates, suggesting that its effect on glutaminase in these models is minimal. Yet both CB-839 and JHU-083 impair tumor growth, at least modestly. These observations suggest that it may be necessary to impair amidotransferases and glutaminase concurrently in order to maximize the therapeutic benefit in ccRCC. Developing a strategy with an acceptable therapeutic index would likely benefit from defining which amidotransferase activities are most important for ccRCC growth.

## Materials and Methods

### Patient-derived xenografts (Tumorgrafts)

All mouse experiments were approved by the UT Southwestern IACUC. All tumorgrafts used in this study were pre-established patient-derived xenografts. For tumor passaging and metabolomics analysis, tumorgrafts were passaged in the left kidney of 4-8 weeks old male/female NOD-SCID mice, as described previously (*29-31*). Once the tumor was palpable and close to ∼ 10mm in diameter, mice were euthanized to collect tumors in ice-cold HBSS buffer with 1% Pen/Strep. For tumor passaging, tumor tissues were dissected aseptically into ∼ 8 mm^3^ of tissues. They were freshly implanted into the left kidney of male or female mice, and the mice were monitored periodically for tumor growth. For subcutaneous tumor studies, resected orthotopic tumors were dissected into small pieces (∼64 mm^3^) and surgically implanted subcutaneously in the right hind region of the mice. Experiments with both orthotopic and subcutaneous tumors were conducted by surgically implanting tumors simultaneously in the left kidney and right hind region of the mice.

### Metabolomics

For tissue metabolomics, mice were sacrificed, and tissues were collected within 3 minutes in liquid nitrogen to minimize metabolite degradation. Tumor and kidney tissues from mice were dissected on dry ice, and ∼50mg of the tissues were collected in 1ml ice-cold 80% MeOH solution in HPLC grade water. Tissues were homogenized and subjected to three freeze-thaw cycles between liquid nitrogen and a 37 °C water-bath. Supernatants were collected by centrifuging the homogenized solution at 10,000 rpm for 10 minutes to remove tissue debris, then the supernatants were evaporated overnight in a SpeedVac concentrator (Thermo Savant). Dried samples were reconstituted in 100□l of 0.03% formic acid in HPLC grade water, vortexed, centrifuged, and the supernatant was transferred in LC/MS glass vials containing inserts. As described previously (*58*), LC/MS was performed using AB QTRAP 5500 liquid chromatography/triple quadrupole mass spectrometer (Applied Biosystems SCIEX, Foster City, CA). Chromatograms and peak area of each metabolite were reviewed and integrated using MultiQuant software version 2.1 (Applied Biosystems SCIEX, Foster City, CA). Each metabolite’s peak area was selected from either positive or negative modes depending on previously run standards. Peak areas were corrected to blank samples and then normalized to the total ion count (TIC) of that sample to correct for variation introduced by sample handling and instrumentation. TIC-normalized data were log-transformed and median normalized for multivariate analyses using Metaboanalyst 4.0 (*59*) (http://www.metaboanalyst.ca). Additional statistical analyses were performed using one-way ANOVA and Student’s t-test in R software.

### Isotope infusions

Isotope infusion experiments were conducted when tumor size was between 10-50 mm diameter for orthotopic tumors and 0.5-1cm diameter for subcutaneous tumors. When the tumors reached these sizes, the mice were fasted for 16 hours and infused in the morning with glucose for 3 hours or glutamine for 5 hours. The mice were anesthetized on a heating pad, and a catheter (connected to the infusion solution and pump) was inserted in the lateral tail vein. For [U-^13^C]glucose infusions, we prepared a 750 μl of saline solution containing 2.48 grams of isotope-labeled glucose per kg of body weight. An initial bolus of 125 μl/min was delivered under anesthesia for 1 min, followed by 2.5 μl/min of continuous infusion for 3 hours. For glutamine infusions, we prepared 1500 μl of a saline solution containing 50 U/ml of Heparin with 1.725 grams of isotope-labeled glutamine per kg of body weight. An initial bolus of 150 μl/min was delivered under anesthesia for 1 min, followed by 2.5 μl/min of continuous infusion for 5 hours. Mice were kept under anesthesia throughout the experiment, and blood samples were collected at regular intervals via retro-orbital puncture using microcapillary tubes. At the end of the infusion, mice were euthanized, and tumors and other organs were harvested and snap-frozen in liquid nitrogen. Endpoint blood samples were collected from the heart with an insulin syringe.

### Gas-chromatography mass-spectrometry (GC-MS)

Blood samples from all infusion experiments were kept on ice until the last sample was collected. Plasma was collected from blood samples through centrifugation at 5000 rpm for 5 minutes at 4 ºC. While 10μl of plasma was directly resuspended in ice-cold 80% MeOH in water, tissues samples of about 5-20mg were homogenized in 80% MeOH in water and subjected to three cycles of free-thaw in liquid nitrogen and a 37 °C water-bath. Tissue and plasma extracts were centrifuged at 10,000 g for 10 min at 4 ºC to separate macromolecules and cell-debris. The supernatants were vacuum dried over-night. The dried samples were reconstituted in 40 μl of pyridine containing methoxyamine (10mg/ml). The samples were vortexed, transferred to GC/MS glass vials with inserts, and heated for 10 minutes at 70 °C. Next, we added 80 μl of the derivatization agent (N-(*tert*-butyldimethylsilyl)-N-methyltrifluoroacetamide (MTBSTFA)) and heated the samples for 1 hour at 70 °C on a heating block. Samples were then loaded into an autoinjector, and 1μl of each sample was injected for analysis. Samples were run on either Agilent 6890 gas chromatograph coupled to an Agilent 5973N or Agilent 7890 gas chromatography coupled to 5975C Mass Selective Detector. The resulting chromatogram and mass isotopologue distributions were analyzed in Agilent MSD ChemStation Data-Analysis software. The area under the curve for mass isotopologues was corrected for natural abundance using in-house methods.

### Liquid-chromatography and mass spectrometry (LC-MS/MS)

For the quantification of NAD, NADH^+^, alpha-ketoglutarate and the analysis of metabolites labeled from [amide-^15^N]glutamine, or [U-^13^C,^15^N]glutamine, we used HILIC column-based separation of metabolites on a Vanquish UHPLC coupled to a QExactive HF-X hybrid quadrupole orbitrap high-resolution mass spectrometer (HRMS) from Thermo Fisher Scientific (Bremen, Germany), as described previously(*60*). Briefly, ∼10-20mg tissue samples were extracted in 80% acetonitrile, homogenized, subjected to 3 freeze-thaw cycles as described above, and cleared of macromolecules by centrifugation. Protein concentration was determined in each supernatant using the BCA Assay, and equivalent aliquots of protein-normalized samples were injected into the instrument. The chromatogram was analyzed as described previously ^52^ using TraceFinder Software from Thermo Fisher Scientific. The ^13^C and ^15^N labeled data were corrected for natural abundance using IsoCorrectoR (*61*).

For targeted GSH quantification, metabolites were extracted from tissues homogenized in 80% acetonitrile solution containing N-ethylmaleimide (0.3 M), followed with 3 freeze-thaw. The samples were centrifuged, and the supernatant was analyzed with a SCIEX QTRAP 5500 liquid chromatography/triple quadrupole mass spectrometer. We performed separation of metabolites on a SeQuant® zic®-pHILIC polymeric HPLC column (150×2 mm) in a Nexera Ultra-High-Performance Liquid Chromatography system (Shimadzu Corporation). Mobile phase solvents were 10 mM ammonium acetate aqueous (pH 9.8 adjusted with ammonia water (A), and pure acetonitrile (B). The gradient elution was: 0– 18 min, linear gradient 90–55% B and 18–20 min, linear gradient 55-30% B, 20-25 min, 30% B, 25-27 min, linear gradient 30-90% B, then the column was reconditioned for 6 min using 90% B. The flow rate was 0.2 ml/min and the column was operated at 40 °C. Multiple reaction monitoring (MRM) was used to check N-ethylmaleimide derivatized GSH, 433/304 (M+0, CE: 19V), 438/304 (M+5, CE: 19V).

### Measurement of D-2HG and L-2HG in tissues

Metabolites were extracted from the tissues samples with 80% methanol-water solution, and the resulting supernatant was dried in a SpeedVac. Dried pellet from unlabeled samples was mixed with [U^13^C]-D/L-2HG (internal standard for unlabeled samples, Cambridge isotope laboratories, 10 nG in 10 μl acetonitrile). The mixture (or pellet from labeled samples) was dissolved in a diacetyl-L-tartaric anhydride(DATAN) solution (90 μl, 50 mG/ml in freshly mixed 80% acetonitrile/20% acetic acid, DATAN, Acros Organics). The solution was sonicated, warmed up to 75 °C and kept for 30 min, cooled to room temperature, and then centrifuged again to collect the supernatant. The supernatant was again dried with a SpeedVac, and the pellet was reconstituted into 1.5 mM ammonium formate aqueous solution with 10% acetonitrile (100 μl). For [U-^13^C]glutamine tracing samples, derivatized [U^13^C]-D/L-2HG was used as standard spiking to identify the derivatized peaks. LC/MS analysis was performed on an AB Sciex 5500 QTRAP liquid chromatography/mass spectrometer (Applied Biosystems SCIEX) equipped with a triple quadrupole/iontrap mass spectrometer with electrospray ionization interface, and controlled by AB Sciex Analyst 1.6.1 Software. Waters Acquity UPLC HSS T3 column (150 × 2.1 mM, 1.8 μM) column was used for separation. Solvents for the mobile phase were 1.5 mM ammonium formate aqueous (pH 3.6 adjusted with formic acid (A), and pure acetonitrile (B). The gradient elution was: 0–12 min, linear gradient 1–8% B and 12– 15 min, 99% B, then the column was washed with 99% B for 5 min before reconditioning it for 3 min using 1% B. The flow rate was 0.25 ml/min and the column was operated at 35 °C. Multiple reaction monitoring (MRM) was used to check 2-hydroxyglutarate-diacetyl tartrate derivatives: 363/147 (M+0, CE: −14V); 368/152 (M+5, CE: −14V). This method was modified and adapted from previously published work (*62*).

### ^13^C Nuclear Magnetic Resonance Spectroscopy

Kidney (43 – 107 mg) and tumor (220 – 462 mg) samples were subjected to extraction in ice cold acetonitrile – isopropanol – water (ACN/IPA/water) by adding the solvent to pre-weighed tissue samples. To this suspension, zirconium beads were added, and the samples were homogenized using a Fastprep-24 (MP Biomedicals, California, USA). Homogenization was carried out in a “20 seconds ON – 5 minutes OFF” cycle to avoid overheating extracts and repeated at least four times. The Homogenates were centrifuged at 10,000xg for 30 minutes at 4°C and the supernatant was lyophilized at room temperature in a Speedvac system (Thermo Scientific, Waltham, MA). The dried sample was dissolved in a 1:1 acetonitrile-water mixture, centrifuged at 10,000xg for 30 minutes at 4°C and lyophilized again.

Samples were prepared for NMR analysis by dissolving the powder in 54 μl of 50 mM sodium phosphate buffer in D_2_O containing 2 mM ethylenediaminetetraacetic acid (EDTA). To this solution, 6 μl of an internal standard containing 0.02%(w/v) NaN_3_ and 0.5 mM deuterated sodium 3 – trimethylsilyl – 1 – propanesulphonate (d6-DSS) was added. The solution was vortexed thoroughly, centrifuged at 10,000xg for 5 minutes and 54 μl of the supernatant was loaded into a 1.5 mm NMR tube. All NMR spectra were acquired using a 14.1 T NMR magnet equipped with a home-built superconducting (HTS) probe (*63*). Parameters included an acquisition time of 1.5 s, a spectral width of 240 ppm, and ^1^H decoupling field strength of 4800 Hz. The spectra were processed (zero-filled to 131,072 points, 0.8 Hz exponential line broadening, and polynomial or spline baseline correction as necessary) and referenced by setting the singlet resonance of taurine (N-C1) to 48.4 ppm. Peaks of interest were fitted to a mixed Gaussian/Lorentzian shape and area under the peaks were obtained.

### L-[5-^11^C]glutamine PET/CT and MRI

A 3.5-inch-long catheter of 0.38 mm inner diameter fitted with a 27-gauge needle was inserted into the tail of a tumor-bearing mouse for intravenous injection of L-[5-^11^C]-glutamine (∼ 5 MBq per mouse). Dynamic PET data acquisition was performed on a Mediso NanoScan PET/CT System (Mediso, USA) immediately after the injection for up to 40 minutes, followed by a CT scan with 720 projections. The images were reconstructed using the manufacturer’s software and quantitatively analyzed by Inveon Research Workplace (Siemens, USA). MR Imaging (MRI) was performed on a 1Tesla Desktop magnetic resonance (MR) scanner (M2 Compact, Aspect Imaging, Shoham, Israel) using a mouse volume coil. T2-weighted imaging was performed with a fast spin-echo (FSE; TR/TE = 2500/80 ms) sequence with the mouse in a prone position.

### Cell lines derived from XP258 tumorgrafts

Orthotopic tumors were dissected and collected in ice-cold 20 ml transport media (MEM, 50 μg/ml gentamycin, 2.5 μg/ml fungizone, and 1X pen/strep). The tissues were washed three times in 15 ml of transport media and minced into small pieces using sterile Swan-Motron blades. Minced tissues were resuspended in a 15 ml enzyme mix (transport media, collagenase, hyaluronidase, and DNase IV and incubated for 1 to 2 hours in 5% CO_2_ in a 37 °C incubator. Enzyme-digested tissue samples were filtered to separate cells from debris and larger tissue pieces. Freshly obtained cells were cultured in high glucose DMEM supplemented with 10% FBS, 1% Pen/Strep, 1X non-essential amino acids, 0.8 mg/ml hydrocortisone and 10 μg/ml EGF.

### Knockout of IDH1 and IDH2 using CRISPR-Cas9

IDH1 and IDH2 knockout pools were generated using the CRISPR-Cas9 system, as described previously (*64*). Briefly, IDH1 sgRNA (ACGTGGAATTGGATCTACAT) and IDH2 sgRNA (ATGAGATGACCCGTATTATC) were cloned into lentiCRISPR V2 system following a published protocol(*65*). These vectors were transfected into HEK239 cells, and viral particles were collected from spent media at 48 hours and 72 hours. The tumorgraft-derived XP258 cell line was transduced with viral particles, then selected in 2βg/ml of puromycin to generate pools of modified cells. IDH1/2 double KO cells were generated by transducing IDH2 KO cells with viral particles carrying the IDH1 sgRNA.

### Immunoblotting

Cells and tissues were homogenized in RIPA buffer containing protease and phosphatase inhibitors. Supernatants were collected by centrifugation at 14,000 RPM for 20 minutes at 4 °C. Protein concentrations were determined using the BCA protein assay. Protein lysates were resolved in SDS-PAGE and transferred to methanol-activated nitrocellulose membranes. The membrane was first blocked with 5% milk and then incubated overnight with primary antibodies GLS (ThermoFisher, Cat# PA5-40135), IDH1 (Cell Signaling, Cat# 8137), IDH2 (Abcam, ab55271), β-Actin or GAPDH from cell signaling.

### CB-839 and JHU-083 treatment

Calithera provided the solution for CB-839 and vehicle for in vivo experiments. 200 mg/kg of CB-839 or vehicle in 200μl of the solution was administered orally every 12 hours for 21-22 days, as described previously (*46*). Barbara Slusher provided JHU-083, and we used a stock solution of 1.83 mg/kg in 50mM HEPES-buffered saline. Mice were dosed with JHU-083 and vehicle via IP injection according to a regimen of 5 days on and 2 days off. Tumor growth and body weight were monitored regularly. Tumor volume was measured using electronic calipers and calculated using the formula Length x (Width^2^)/2. Infusion of [U-^13^C]glutamine and metabolomics were conducted at the end of 21 days of the CB-839 treatment. Infusion with [Amide-^15^N]glutamine, [U-^13^C,^15^N]glutamine, and [U-^13^C]glucose were conducted after 5-7 doses of the drugs (JHU-083 or CB-839) and vehicle.

### Isotope-tracing in vitro

One million cells were plated overnight for each experiment. For isotope-tracing experiments, cells were washed with PBS and then fed with medium containing 100% isotope-labeled compound with 10% dialyzed FBS. [U-^13^C]glucose and [U-^13^C]glutamine labeling was performed over 3-4 hours. After the experiment, the cells were washed once in ice-cold saline solution and 1ml of 80% methanol solution was added. The cells were collected by scraping and subjected to a freeze-thaw cycle followed by centrifugation to remove debris. The supernatant was vacuum dried overnight and reconstituted for GC-MS analysis as described above.

### Statistical analysis

Metaboanalyst 4.0(*59*) and R-software (http://www.metaboanalyst.ca) were used to conduct statistical analysis of the metabolomics data. R software and PRISM were used to conduct one-way ANOVA and Student’s t-test, respectively. Data visualization used both PRISM and R Software. Descriptions of individual statistical analyses can be found in the Fig. legends.

## Acknowledgments

We are grateful to the patients who provided tissue for the tumorgrafts we studied; to members of the DeBerardinis laboratory for constructive comments on the project and the manuscript; and to Calithera for providing CB-839.

## Funding

H.H.M.I. Investigator Program (RJD),

NCI Outstanding Investigator Award (R35CA22044901 to RJD),

Kidney Cancer SPORE grant (P50CA19651602 to JB),

2019 Young Investigator Award, Kidney Cancer Association (AKK)

### Author contributions

Conceptualization: AKK, RJD

Methodology: AKK, LKB, AT, MR, CYW, XL, KA, VTT, FS, QND, CY, TR, CS, AR, BF, PP, HV, FC, MM, BS, TPM, PK, JB

Investigation: AKK, MR, CYW, LGZ, FC

Visualization: AKK, MR, CYW, QND, XS, RJD

Supervision: RJD

Writing—original draft: AKK, RJD

Writing—review & editing: AKK, MR, XS, MM, JB, RJD

### Data and materials availability

The data can be provided by RJD pending scientific review and a completed material transfer agreement. Requests for the data should be submitted to the corresponding author.

