## Supplemental Data for "In-vivo characterization of glutamine metabolism identifies therapeutic targets in clear cell renal cell carcinoma"

#### This PDF file includes:

Figs. S1 to S8
Table S1

#### Other Supplementary Materials for this manuscript include the following:

Data S1, S2

Figure S1

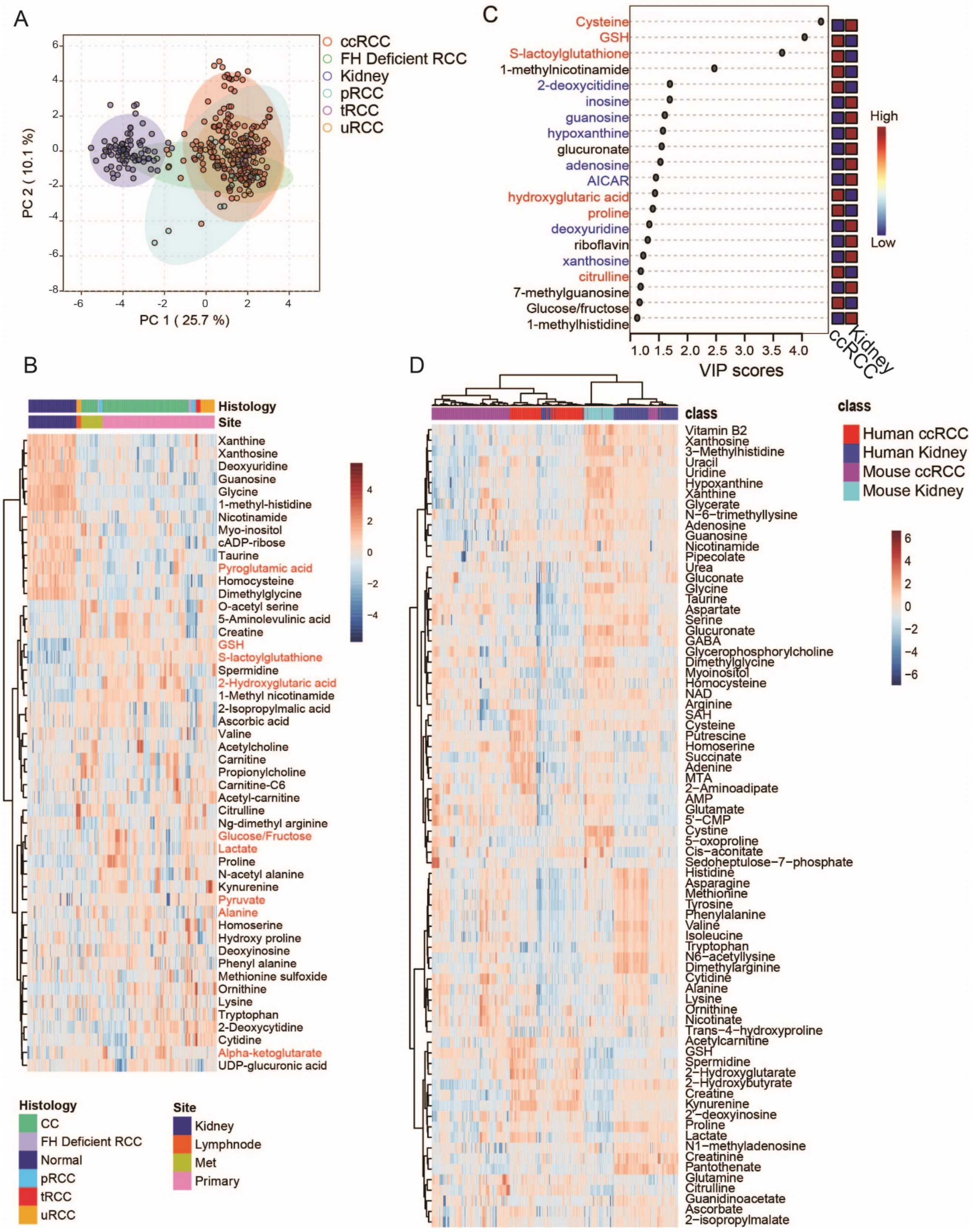

A
ccRCC
FH Deficient RCC
Kidney
pRCC
tRCC
uRCC
PC 2 ( 10.1 %)
PC 1 ( 25.7 %)
B
Histology
Site
Xanthine
Xanthosine
Deoxyuridine
Guanosine
Glycine
1-methyl-histidine
Nicotinamide
Myo-inositol
cADP-ribose
Taurine
Pyroglutamic acid
Homocysteine
Dimethylglycine
O-acetyl serine
5-Aminolevulinic acid
Creatine
GSH
S-lactoylglutathione
Spermidine
2-Hydroxyglutaric acid
1-Methyl nicotinamide
2-Isopropylmalic acid
Ascorbic acid
Valine
Acetylcholine
Carnitine
Propionylcholine
Carnitine-C6
Acetyl-carnitine
Citrulline
Ng-dimethyl arginine
Glucose/Fructose
Lactate
Proline
N-acetyl alanine
Kynurenine
Pyruvate
Alanine
Homoserine
Hydroxy proline
Deoxyinosine
Phenyl alanine
Methionine sulfoxide
Ornithine
Lysine
Tryptophan
2-Deoxycytidine
Cytidine
Alpha-ketoglutarate
UDP-glucuronic acid
Histology
CC
FH Deficient RCC
Normal
pRCC
tRCC
uRCC
Site
Kidney
Lymphnode
Met
Primary
C
Cysteine
GSH
S-lactoylglutathione
1-methylnicotinamide
2-deoxycitidine
inosine
guanosine
hypoxanthine
glucuronate
adenosine
AICAR
hydroxyglutaric acid
proline
deoxyuridine
riboflavin
xanthosine
citrulline
7-methylguanosine
Glucose/fructose
1-methylhistidine
VIP scores
ccRCC
Kidney
High
Low
D
class
Human ccRCC
Human Kidney
Mouse ccRCC
Mouse Kidney
Vitamin B2
Xanthosine
3−Methylhistidine
Uracil
Uridine
Hypoxanthine
Xanthine
Glycerate
N−6−trimethyllysine
Adenosine
Guanosine
Nicotinamide
Pipecolate
Urea
Gluconate
Glycine
Taurine
Aspartate
Serine
Glucuronate
GABA
Glycerophosphorylcholine
Dimethylglycine
Myoinositol
Homocysteine
NAD
Arginine
SAH
Cysteine
Putrescine
Homoserine
Succinate
Adenine
MTA
2−Aminoadipate
AMP
Glutamate
5'−CMP
Cystine
5−oxoproline
Cis−aconitate
Sedoheptulose−7−phosphate
Histidine
Asparagine
Methionine
Tyrosine
Phenylalanine
Valine
Isoleucine
Tryptophan
N6−acetyllysine
Dimethylarginine
Cytidine
Alanine
Lysine
Ornithine
Nicotinate
Trans−4−hydroxyproline
Acetylcarnitine
GSH
Spermidine
2−Hydroxyglutarate
2−Hydroxybutyrate
Creatine
Kynurenine
2'−deoxyinosine
Proline
Lactate
N1−methyladenosine
Creatinine
Pantothenate
Glutamine
Citrulline
Guanidinoacetate
Ascorbate
2−isopropylmalate

**Figure S1. Metabolic altertaions in RCC tumorgrafts and similarlity between tumorgrafts and published human studies.**

(A). Principal component analysis (PCA) plot of metabolomics data generated from 28 distinct RCC tumorgrafts passaged in NOD-SCID mice. Metabolomics data were log-transformed and median normalized to assess metabolic differences between normal kidney tissues and RCC subtypes using Metaboanalyst.

(B). Heatmap of the top 50 differential metabolites between RCC subtypes and kidney tissues. One-way ANOVA was used to assess the significance of the metabolites. Metabolites of central carbon metabolism and glutathione metabolism are shown in red fonts. The heatmap was generated using the default parameter in Metaboanalyst.

(C). Variable importance in projection (VIP) plot of the top 20 metabolites differing between ccRCC and kidney using partial least squares-discriminant analysis (PLS-DA) in Metaboanalyst.

(D). Heatmap of 76 common metabolites between tumorgrafts and published data from Hakimi et al. Raw data were log-transformed and median-normalized to combine and generate the heatmap using the default parameter in Metaboanalyst.

Figure S2

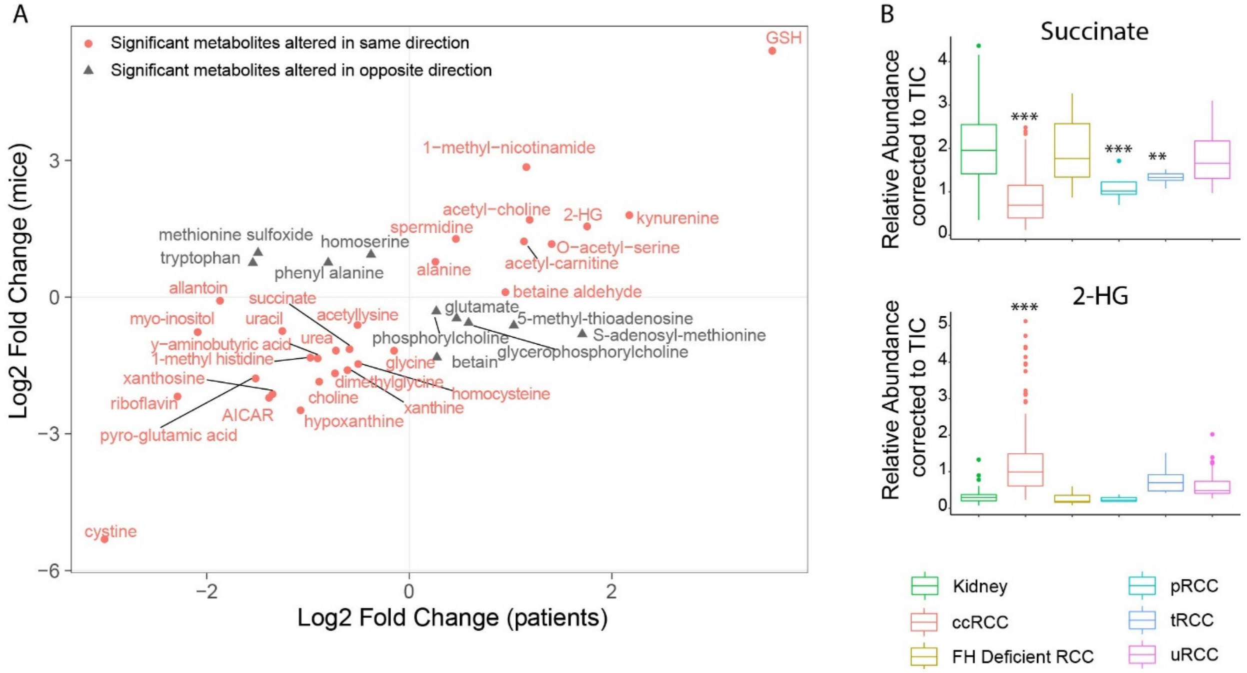


**Figure S2. Correlation of metabolic features between ccRCC tumorgrafts and human studies and metabolic features of RCC tumorgrafts.**

(A). Correlation plot of the metabolites altered in both ccRCC tumorgrafts and published data from Zhang et al. In both studies, differential metabolites between tumor and kidney tissues were calculated using Student's t-test and corrected for FDR (Q value<0.05). Fold change was used as a surrogate for z-score to generate the correlation plot using ggplot in R. Positive Log2 fold change indicate metabolite elevation and negative Log2 fold change indicate metabolite depletion in tumors compared to kidney. Metabolites in red are altered in the same direction in tumorgrafts and patients, whereas metabolites in gray are altered in the opposite direction in tumorgrafts and patients.

(B). Boxplot of the relative abundance of succinate and 2-hydroxyglutarate in RCC tumorgrafts of different histological types, including ccRCC, FH deficient RCC, papillary RCC (pRCC), translocation RCC (tRCC), and unclassified RCC (uRCC). One-Way ANOVA coupled with pairwise t-test in R software was used to calculate the significance of the data and P values were FDR corrected. P values: ***<0.001, **<0.01, *<0.05

Figure S3

A

| Tumorgraft | Mutations | Histology | Previous treatment (patients) |
|---|---|---|---|
| XP258 | VHL | Clear cell with sarcomatoid | None |
| XP296 | VHL | Clear cell with sarcomatoid | Everolimus |
| XP374 | VHL, SETD2, TP53 | Clear cell with sarcomatoid | None |
| XP955 | VHL | Clear cell | None |
| XP490 | VHL, TP53 | Clear cell with sarcomatoid | None |

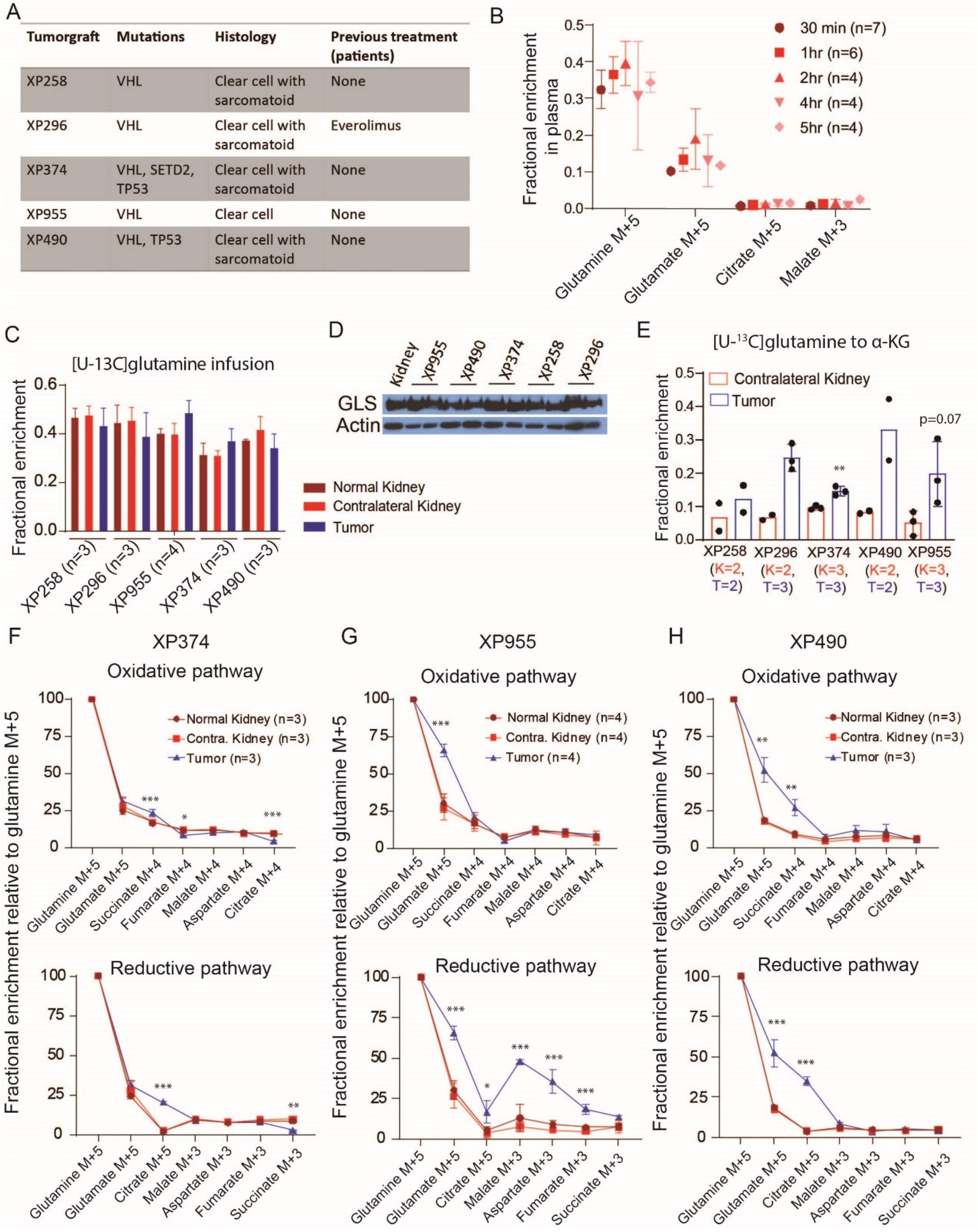


**Figure S3. Characterization of $^{13}$C enrichment in ccRCC tumorgrafts infused with [U-$^{13}$C]glutamine.**

(A). Table showing genetic and histological characteristics of ccRCC tumorgrafts used in this study. We used five distinct *VHL*-mutant ccRCC tumorgrafts. Four tumorgrafts were derived from treatment-naïve patients and one was derived from a patient previously treated with everolimus. Mutations in tumors from both patient and tumorgraft tissues were confirmed using whole-exome sequencing.

(B). Fractional enrichment of glutamine M+5, glutamate M+5, citrate M+5, and malate M+3 in plasma collected at different time points from mice infused with [U-$^{13}$C]glutamine. (C). Fractional enrichment of $^{13}$C-glutamine in the tumor, normal kidney and contralateral kidney collected from mice infused with [U-$^{13}$C]glutamine.

(D). Western blot showing glutaminase (GLS) expression relative to β-actin in contralateral kidney and tumor tissues collected from mice bearing orthotopic ccRCC tumorgrafts.

(E). Fractional enrichment of $^{13}$C labeled α-ketoglutarate (α-KG) M+5 in contralateral kidney and tumorgrafts collected from mice infused with [U-$^{13}$C]glutamine. The number of independent tissues used is displayed as K=n for the kidney and T=n for the tumorgraft. One-way ANOVA was used to determine the statistical significance.

(F). Percentage enrichment of the TCA cycle intermediates relative to [U-$^{13}$C]glutamine in XP374 tumorgraft, adjacent benign (normal kidney) and contralateral kidney. The top panel shows $^{13}$C labeling via the oxidative pathway and the bottom panel shows $^{13}$C labeling via the reductive pathway. The experiment was conducted in a minimum of 3 mice per tumorgraft model. One-way ANOVA was used to assess the statistical significance of $^{13}$C enrichment between tissues.

(G). Same as in E, but for XP955 tumorgrafts.

(H). Same as in E, but for XP490 tumorgrafts. P values: ***<0.001, **<0.01, *<0.05

Figure S4

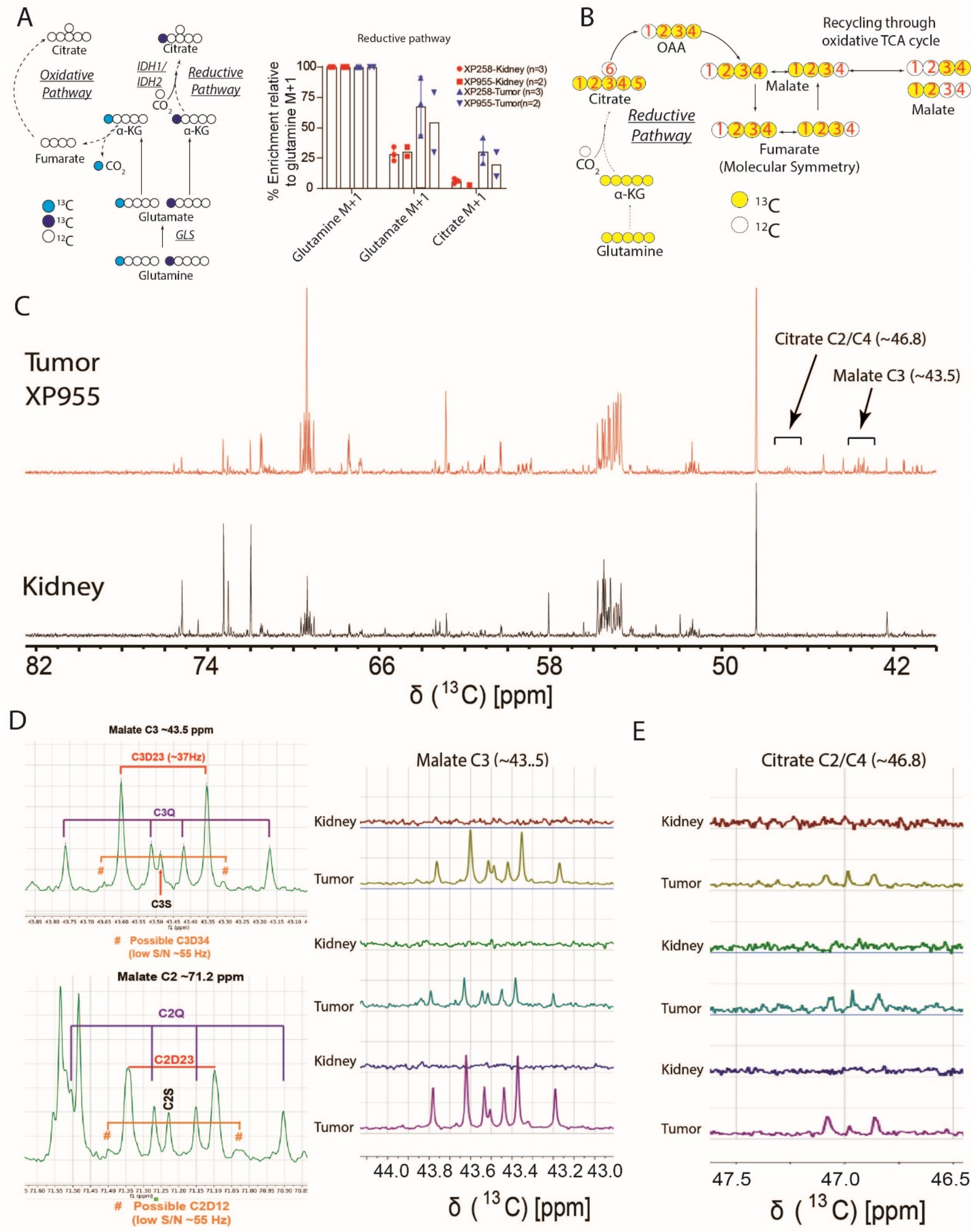
A
Citrate
Oxidative Pathway
Fumarate
CO2
IDH1/ IDH2
Reductive Pathway
α-KG
Glutamate
GLS
Glutamine
13C
13C
12C
Reductive pathway
% Enrichment relative to glutamine M+1
XP258-Kidney (n=3)
XP955-Kidney (n=2)
XP258-Tumor (n=3)
XP955-Tumor(n=2)
Glutamine M+1
Glutamate M+1
Citrate M+1
B
OAA
Recycling through oxidative TCA cycle
Malate
Fumarate
(Molecular Symmetry)
Citrate
Reductive Pathway
CO2
α-KG
Glutamine
13C
12C
C
Tumor XP955
Kidney
Citrate C2/C4 (~46.8)
Malate C3 (~43.5)
δ (13C) [ppm]
D
Malate C3 ~43.5 ppm
C3D23 (~37Hz)
C3Q
C3S
### Possible C3D34 (low S/N ~55 Hz)
Malate C2 ~71.2 ppm
C2Q
C2D23
C2S
### Possible C2D12 (low S/N ~55 Hz)
Malate C3 (~43..5)
Kidney
Tumor
δ (13C) [ppm]
E
Citrate C2/C4 (~46.8)
Kidney
Tumor
δ (13C) [ppm]

**Figure S4**. **GC-MS and NMR in ccRCC tumorgrafts infused with [U-$^{13}$C]glutamine.**

(A). Left panel shows the schematic representation of $^{13}$C labeling from [1-$^{13}$C]glutamine. Citrate is labeled via the reductive pathway. The light blue-filled circles show $^{13}$C labeling via the oxidative pathway, and the dark blue-filled circles show $^{13}$C labeling via the reductive pathway. The right panel shows the percentage enrichment of $^{13}$C in glutamine M+1, glutamate M+1, and citrate M+1 relative to [1-$^{13}$C]glutamine in tumor and contralateral kidney from two tumorgraft models (XP258, n=3; XP955, n=2).

(B). Schematic of $^{13}$C positions in TCA cycle intermediates via the reductive pathway. The yellow circles represent $^{13}$C, and the numbers in the circles represent the position of the $^{13}$C. Although reductive carboxylation initially generates malate and fumarate labeled with $^{13}$C at positions 2, 3 and 4 (234), 123 labeling equilibrates with 234 labeling due to molecular symmetry of fumarate. Labeling at 12 and 34 in malate arise from processing along the oxidative pathway.

(C). $^{13}$C NMR spectra of an XP955 tumorgraft (red) and contralateral kidney (black) collected from mice infused with [U-$^{13}$C]glutamine. The spectra corresponding to citrate C2/C4 (~46.8 ppm) and malate C3 (~43.5 ppm) are highlighted. These spectra are representative of 3 biological experiments.

(D). In the left panel, malate C3 and C2 spectra are shown, highlighting peaks for singlets (S: C3S or C2S), doublets (D: C3D23 and C3D34; or C2D12 and C2D23), and quartets (Q: C3Q or C2Q). The predicted ppm for C3D34 and C2D12 multiplets generated via the oxidative TCA cycle are highlighted in orange. The right panel shows spectra for malate C3 (~43.5 ppm) across all samples (n=3 tumors and 3 kidneys).

(E). Spectra for citrate C2/C4 (~46.8 ppm) for all samples (n=3 tumors and 3 kidneys).

Figure S5

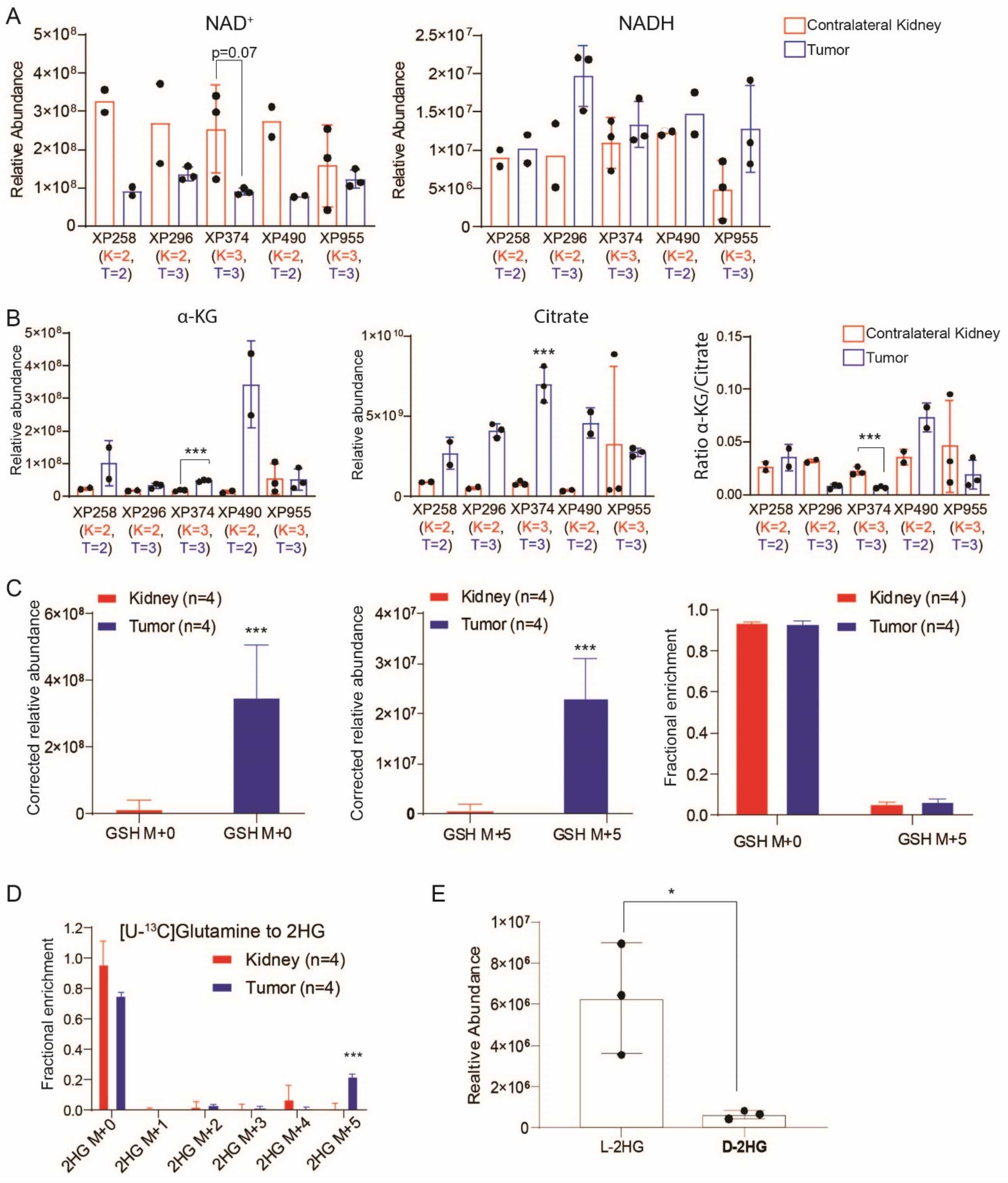
A
NAD+
NADH
Contralateral Kidney
Tumor
Relative Abundance
p=0.07
XP258 (K=2, T=2)
XP296 (K=2, T=3)
XP374 (K=3, T=3)
XP490 (K=2, T=2)
XP955 (K=3, T=3)
B
α-KG
Citrate
Relative abundance
Ratio α-KG/Citrate
***
C
Kidney (n=4)
Tumor (n=4)
Corrected relative abundance
Fractional enrichment
GSH M+0
GSH M+5
D
[U-13C]Glutamine to 2HG
Fractional enrichment
2HG M+0
2HG M+1
2HG M+2
2HG M+3
2HG M+4
2HG M+5
E
Realtive Abundance
L-2HG
D-2HG
*

**Figure S5. Assessment of the relative abundance of $NAD^+$, NADH, citrate, and α-KG, and the fractional enrichment of GSH, and 2HG levels in ccRCC tumorgrafts infused with [U-$^{13}$C]glutamine.**

(A). Relative abundance of $NAD^+$ and NADH in ccRCC tumorgrafts and contralateral kidney, normalized to protein abundance. The number of tissues used for analysis is shown below each plot as K=n for kidney and T=n for tumor. One-way ANOVA was used to determine statistical significance.

(B). Relative abundance of α-KG, citrate, and α-KG/citrate ratio in ccRCC tumorgrafts and contralateral kidneys. The number of tissues used for the analysis is shown below each plot as K=n for kidney and T=n for tumor. One-way ANOVA was used to determine statistical significance.

(C). Corrected abundance and fractional enrichment of M+0 and M+5 $^{13}$C labeled glutathione (GSH) in tumors (XP955) and kidneys of mice infused with [U-$^{13}$C]glutamine. The Student's t-test was used to determine the p-value.

(D). Fractional enrichment of $^{13}$C -abeled 2HG in tumor (XP955) and kidney of the mice infused with [U-$^{13}$C]glutamine. One-way ANOVA was used to determine the p-value

(E). Relative abundance of L-2HG and D-2HG isoforms in ccRCC tumorgrafts (XP296). Student's t-test was used to calculate the p-value.

P values: ***<0.001, **<0.01, *<0.05

Figure S6

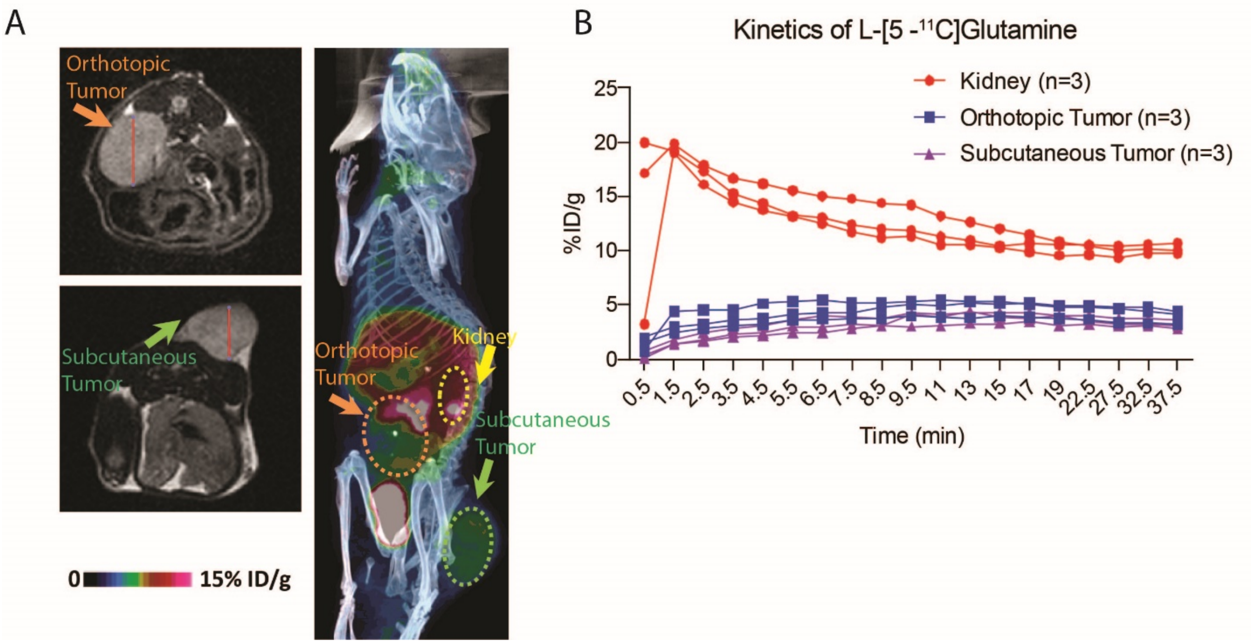


**Figure S6**. **MRI and L-[5-$^{11}$C]glutamine PET/CT analysis in ccRCC tumorgrafts.**

(A). MRI and PET/CT of the mice bearing both orthotopic and subcutaneous XP490 tumorgrafts (n=3 mice). MRI was used to localize the relevant tissues in these imaging planes. PET/CT images show the signal from L-[5-$^{11}$C]glutamine for kidney and tumors. For each mouse, MRI and PET/CT images are placed side-by-side and the location of tumors and kidney are highlighted.

(B). Shows the kinetics of L-[5-$^{11}$C]glutamine as %injected dose per gram of mouse (%ID/g) in the kidney (red), orthotopic tumor (blue), and subcutaneous tumor (purple). Data plotted from n=3 mice.

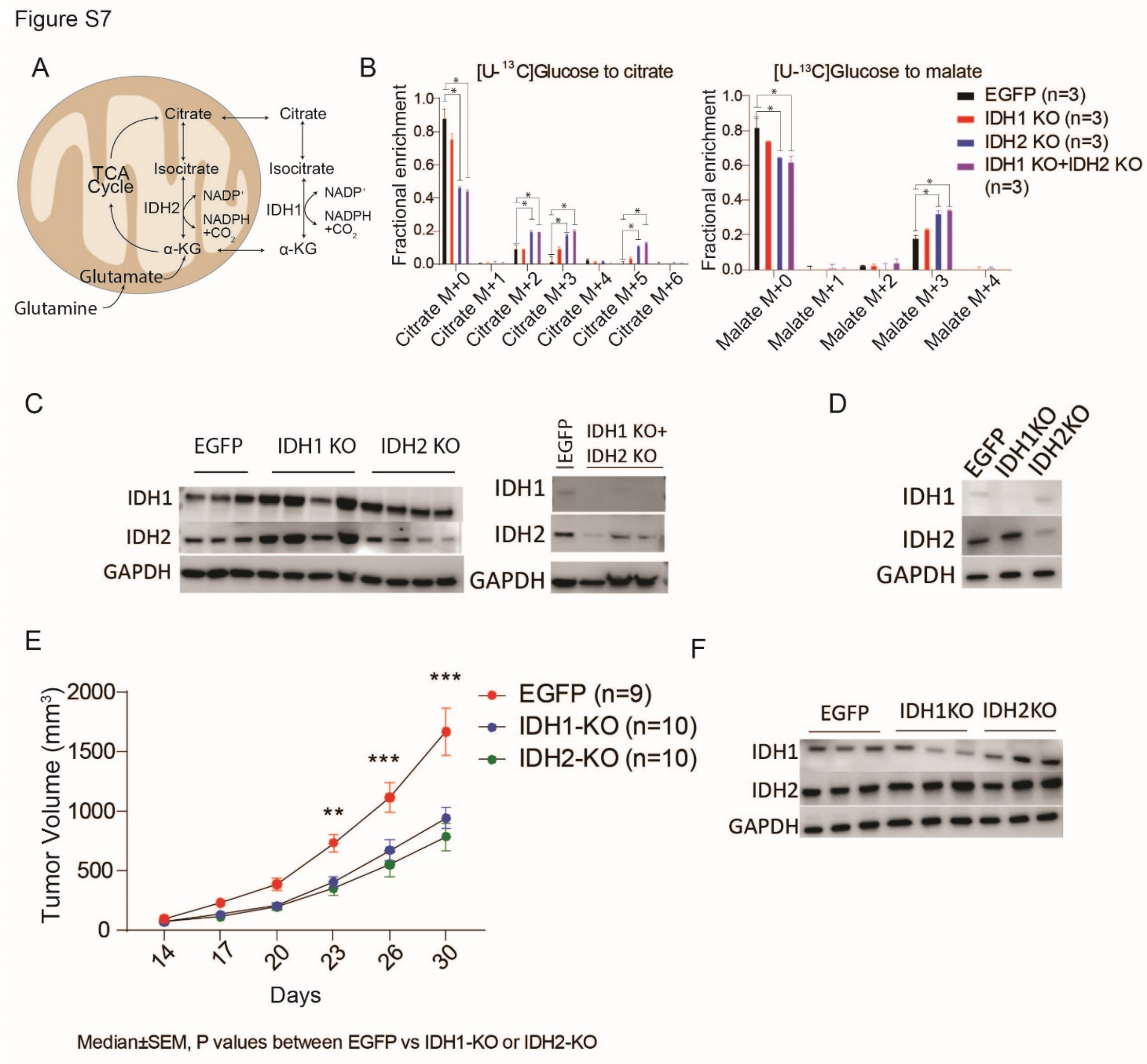


**Figure S7. IDH1 and IDH2 KO rewire substrate oxidation in TCA cycle and reduce tumor growth in ccRCC.**

(A). Schematic of $NADP^+$/NADPH-dependent IDH1 (cytosol) and IDH2 (mitochondria) reactions that interconvert isocitrate and α-KG. Glutamine-derived α-KG is reductively carboxylated by NADPH-dependent IDH1 and IDH2 in cytosol and mitochondria, respectively.

(B). Fractional enrichment of $^{13}$C-labeled isotopologues of citrate and malate in EGFP, IDH1 KO, IDH2 KO, and IDH1 KO +IDH2 KO (double KO) cells labeled with [U-$^{13}$C]glucose for 3 hours. One-way ANOVA was used to assess the statistical significance of $^{13}$C enrichment in isotopologues of citrate and malate among the cell lines (n=3).

(C). Protein expression of IDH1 and IDH2 in tissues collected at the end of the tumor growth study. GAPDH was used as a loading control.

(D). Western blot showing the expression of IDH1 and IDH2 protein in CRISPR-Cas9-generated pools containing either EGFP-targeting gRNA or gRNA targeting IDH1 or IDH2. The parental cells were an independent cell line from the one shown in the main figure, derived by a different lab.

(E). Growth of tumors derived from control cells and IDH1 and IDH2 KO cells (n=5 female and 5 male mice in each group, except 4 female in EGFP) from XP258 cells in panel d. One-way ANOVA was used to assess the statistical significance at each time point of tumor measurement.

(F). Western blot showing the expression of IDH1 and IDH2 in tumors from panel E.

P values: ***<0.001, **<0.01, *<0.05

Figure S8

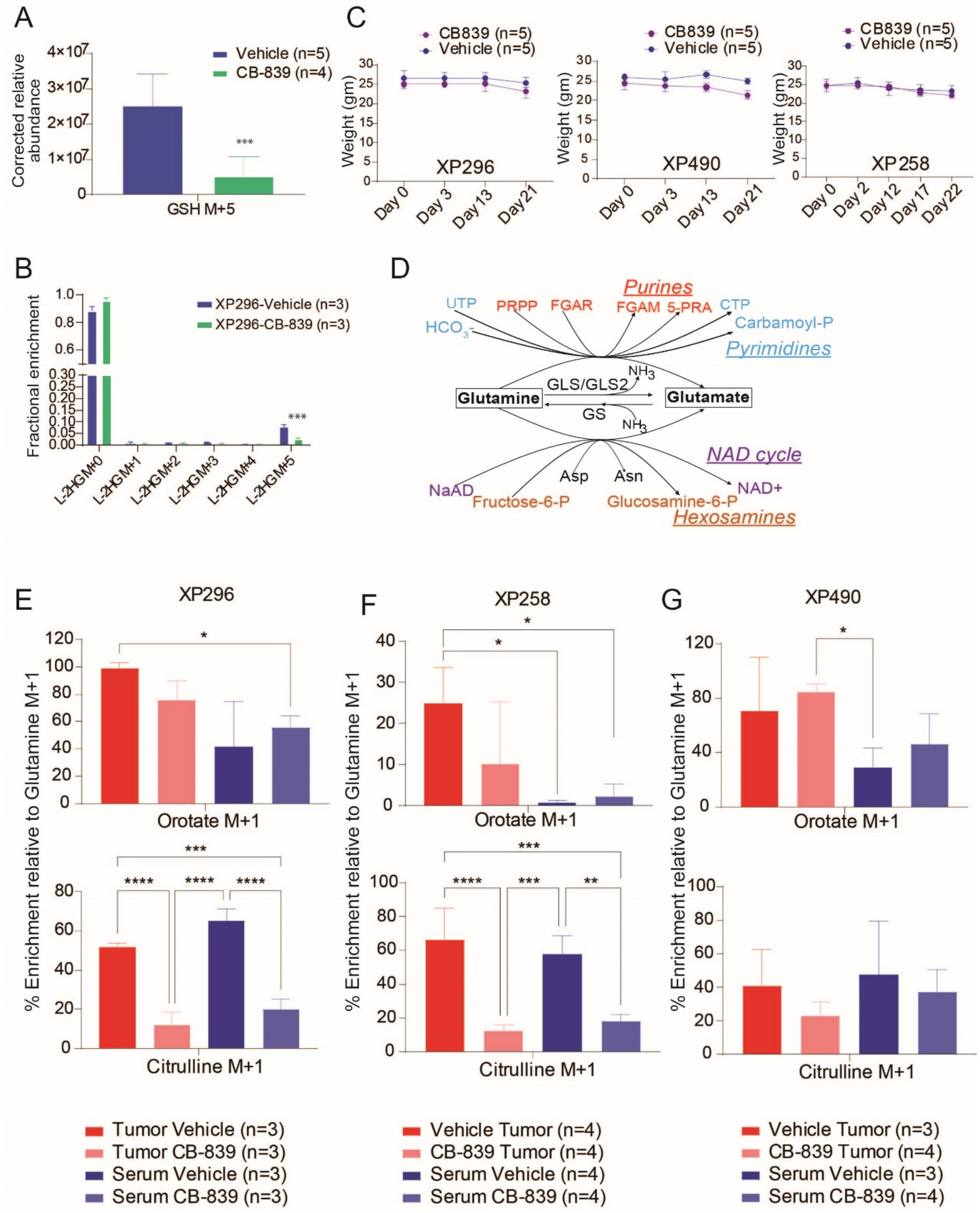


**Figure S8**. **Effects of CB-839 on glutamine-dependent metabolism in ccRCC in vivo.**

(A). Effect of CB-839 on relative abundance of labeled GSH in XP296 tumorgrafts.

(B). Effect of CB-839 on $^{13}C$ labeling in L-2HG (L-2 hydroxyglutarate) in XP296 tumorgrafts. Student's t-test was used to assess the p-value for data represented in panels A and B.

(C). Weight of mice treated with vehicle or CB-839 (200mg/kg twice daily) during tumor growth study.

(D). Conversion of glutamine to glutamate involves glutaminases (GLS, and GLS2) and amidotransferases.

(E). Enrichment relative to glutamine M+1 ([amide-$^{15}N$]glutamine) for orotate and citrulline in plasma and XP296 tumors collected from vehicle or CB-839 treated mice. Mice bearing subcutaneous tumors were treated with 7 doses of either vehicle or CB-839 (200mg/kg twice daily) before infusion with [amide-$^{15}N$]glutamine. Fractional enrichment of each metabolite was corrected to the fractional enrichment of [amide-$^{15}N$]glutamine. P values were calculated using one-way ANOVA.

(F). Same as in E, but for the XP258 tumorgrafts.

(G). Same as in E, but for XP490 tumorgrafts.

P values: ****<0.0001, ***<0.001, **<0.01, *<0.05

Table S1: Clinical characteristics of 28 RCC tumorgrafts.

| **Characteristic** | **N = 28**[1] |
|---|---|
| Previous.Therapies | |
| Everolimus | 1 (3.6%) |
| HD-IL2, Interferon, Bevacizumab | 1 (3.6%) |
| Sunitinib | 3 (11%) |
| Treatment Naïve | 23 (82%) |
| Histological.Subtype | |
| ccRCC | 19 (68%) |
| FH Deficient RCC | 2 (7.1%) |
| pRCC | 2 (7.1%) |
| tRCC | 1 (3.6%) |
| uRCC | 4 (14%) |
| Grade | |
| 3 | 11 (44%) |
| 4 | 13 (52%) |
| High grade | 1 (4.0%) |
| Unknown | 3 |
| Source | |
| Metastasis | 5 (18%) |
| Primary Tumor | 22 (79%) |
| Regional Lymphnode | 1 (3.6%) |
| VHL | |
| Frameshift Indel | 6 (25%) |
| Missense | 5 (21%) |
| Nonsense | 1 (4.2%) |
| Nonstop | 1 (4.2%) |
| WT | 11 (46%) |
| Unknown | 4 |

[1] n (%)
